# Brain network dynamics determine tau presence while regional vulnerability governs tau load in Alzheimer’s disease

**DOI:** 10.1101/2025.04.17.648358

**Authors:** Yu Xiao, Nicola Spotorno, Lijun An, Vincent Bazinet, Justine Y. Hansen, Olof Strandberg, Golia Shafiei, Harry H. Behjat, Thomas Funck, Gemma Salvadó, Erik Stomrud, Ruben Smith, Sebastian Palmqvist, Rik Ossenkoppele, Niklas Mattsson-Carlgren, Nicola Palomero-Gallagher, Alain Dagher, Bratislav Misic, Oskar Hansson, Jacob W. Vogel

## Abstract

In Alzheimer’s disease (AD), tau pathology accumulates gradually throughout the brain, with clinical decline reflecting tau progression. A comprehensive understanding of, first, whether tau propagation is predominantly governed by connectome-based diffusion, regional vulnerability, or an interplay of both, and second, which types of brain connectivity or regional factors best explain tau propagation, remains crucial for advancing our understanding of AD progression. Here, we apply multi-scale, biologically informed disease progression simulations to human data, to disentangle the influence of local mechanisms on global tau progression patterns in AD. We find that whether tau reaches a brain region (presence) and how much tau accumulates there (load) are governed by different mechanisms. Tau presence patterns are highly consistent across the population, and can be largely explained through synaptic spread through white-matter networks and excitatory-inhibitory dynamics. Meanwhile tau load differs across people, and is driven by a combination of synaptic spread and intrinsic or extrinsic regional properties, including regional *β*-amyloid load, *MAPT* gene expression and regional blood flow. Finally, while distinct tau patterns in the population could each be explained by established AD mechanisms, our models highlight a role of distinct brain networks (parietal networks in MTL-sparing AD tau subtype) and neurotransmitter systems (cholinergic system in posterior subtype). Together, this work suggests that network dynamics likely determine the sequence of regional tau progression, while individual-specific tissue-vulnerability factors influence regional tau load.

## Main

Alzheimer’s disease (AD) is the leading cause of dementia worldwide, with its prevalence projected to triple over the next 30 years ^1^. AD is characterized by the accumulation of diffuse extracellular and neuritic amyloid-*β* (A*β*) plaques, alongside intracellular neurofibrillary tangles (NFTs) composed of hyperphosphorylated tau, both of which are fundamental to disease progression ^2,3^. The main hypothetical model of AD progression posits that A*β* accumulation, combined with tau pathology, drives subsequent neurodegeneration and cognitive decline ^4,5^. Compared to A*β*, tau pathology has a stronger predictive value for future neurodegeneration ^6,7^ and a closer association with subsequent cognitive decline ^7–9^, underscoring its importance in the AD pathophysiological cascade. These properties of tau have led to an increased focus on development of tau-targeting drugs ^10–12^. Tau is a complex protein to study, with a wide variety of isoforms and post-translational modifications, which appear to influence properties and location of its accumulation ^13–16^. Understanding the mechanisms underlying tau propagation is therefore crucial for developing more effective tau-targeted interventions, though these mechanisms are not well understood at present.

The accumulation of pathological tau in AD often follows a stereotypical hierarchical spatial pattern, formalized as the Braak staging system ^17–20^. This system delineates the progression of tau from the transentorhinal cortex, to the medial and basal temporal lobes, followed by neocortical associative regions, and eventually the unimodal primary sensory and somatomotor cortices. Braak staging thus provides a semi-quantitative measure of tau progression in AD, indicating the presence of tau tangles across specific brain regions at distinct stages of the disease. However, this approach does not account for the quantitative level of tau accumulation within each region. Tau positron emission tomography (tau-PET) allows quantification of tau *in vivo*, enabling spatially resolved measures of tau pathology across the whole brain ^19–22^. Whole-brain sampling has revealed important spatial characteristics in tau accumulation that complement and extend findings from neuropathology, including a strong link between tau location and clinical symptoms ^8,23–25^ and a spatiotemporal evolution of tau accumulation ^20,26^. Importantly, these studies suggest that individual tau accumulation patterns often diverge from the stereotypical Braak staging system and exhibit substantial variability across individuals ^27–30^.

The mechanisms that underpin tau propagation and the factors that influence this process remain uncertain, though several compelling hypotheses have been proposed ^31–37^. The network spread hypothesis posits that tau pathology propagates from an initial “epicenter” (for example the transentorhinal cortex) along connected brain regions, potentially through synaptic transmission ^38–41^. This view is supported by both *in vivo* human studies ^42–46^, and *in vitro* cell and animal models ^36,47^. However, it is unclear if neuronal connectivity alone is sufficient to predict where tau will distribute. Notably, evidence has emerged for alternative mechanisms of tau spread, including cascading excitotoxic events or altered proteostasis involving A*β* ^48–52^ or glial networks mediating tau phagocytosis and subsequent secretion ^53–55^. Meanwhile, increasing evidence underscores the role of selective regional vulnerability in modulating tau accumulation patterns ^37,56,57^.

Brain regions prone to tau pathology are often characterized by neurofilament-rich excitatory neurons ^56,58–60^, sparse myelination ^61,62^, phylogenic/ontologic immaturity ^63^ and distinct patterns of gene expression ^59,64,65^ or neuroreceptor distribution ^56,66^. This raises the possibility that vulnerability to tau accumulation is an intrinsic property of susceptible neurons, and that its spatiotemporal evolution represents a pathological process traversing along a gradient of vulnerability ^67^. Teasing apart the relative contributions of these different brain properties to the distribution of AD tau pathology will be critical for developing therapies that slow or halt its progress.

Connectome-based disease spreading models offer a mechanistic framework for simulating tau propagation through human brain networks and resolving the mechanisms underlying tau propagation ^67,68^. Studies employing connectome-based spreading models have demonstrated that tau pathology can propagate through human white matter connections measured by diffusion weighted imaging (DWI) ^69–74^. Some studies have more recently modeled regional vulnerability alongside connectome-based spread, building evidence that tau deposition is influenced by factors such as regional microglial activity ^75^ and gene expression patterns ^76^. However, these studies restrict themselves to one or a few predefined factors representing regional vulnerability. A comprehensive understanding of, first, whether tau propagation is predominantly governed by connectome-based spread, regional vulnerability, or an interplay of both, and second, which types of brain connectivity or regional factors best explain tau propagation, remains crucial for advancing our understanding of AD progression.

Here, we present a comprehensive investigation into the mechanisms of tau propagation in AD using tau-PET imaging. We employed the Susceptible-Infected-Removed (SIR) agent-based model ^77,78^, a mechanistic disease progression framework that simulates both brain network spread and cellular tau dynamics at the level of brain regions. This model allows for *in silico* testing of distinct pathophysiological mechanisms by parametrizing how regional factors influence tau synthesis, spread, misfolding, and clearance. We leverage this framework to investigate the interplay between connectome-based spread and regional vulnerability, and to assess how these mechanisms explain population variation in tau accumulation patterns. Our first aim was to understand the overarching mechanisms of tau propagation, determining whether it is primarily driven by connectome-based spread, regional vulnerability, or both. Here, we hypothesized that tau distribution will be explained by a combination of connectome-based spread and regional vulnerability. Second, we test a comprehensive assay of multiscale regional properties of the brain to determine which factors contribute most to tau accumulation patterns. Based on previous work ^59,72,79^, we hypothesized that adding regional Aβ-PET, regional MAPT expression and myelination would improve simulation of tau-PET patterns. However, we also anticipated unexpected measures would emerge that would improve model fit, including metabolic factors and gene expression patterns. Finally, we explore population variation in tau propagation, identifying biological factors that contribute to heterogeneity in tau accumulation patterns. This work provides new insights into the complex dynamics of tau propagation, highlighting factors contributing to this phenomenon.

## Results

This study included 646 A*β*-positive participants from the Swedish BioFINDER-2 study (http://biofinder.se/; NCT03174938). Among them, 219 were identified as cognitively unimpaired (CU), 212 were diagnosed with mild cognitive impairment (MCI), and 215 were diagnosed with AD dementia. Demographic information is shown in Supplementary Table S1.

### Tau load and tau presence index two distinct phenomena

[^18^F]RO948-PET data were used to extract tau standardized uptake value ratios (SUVR) across 66 cortical regions of interest ^80^ for each participant, including the left and right hippocampus and amygdala. To represent tau load, SUVR values were MinMax normalized using a reference group of Aβ-negative, cognitively normal elderly participants, adjusted for choroid plexus binding (Fig. 1b and <u>Methods</u>). The SUVRs for each brain region were then transformed into “tau-positive probabilities” (TPP) using two-component Gaussian mixture models, serving as an index of tau presence (Fig. 1a and <u>Methods</u>). Unlike SUVR, which quantifies the level of tau, TPP reflects a confidence estimate as to whether tau pathology has reached a given region, irrespective of the total amount, thereby capturing the pattern of tau presence rather than its magnitude. When averaged across a population (Fig. 1c,1d), TPP represents the proportion of individuals who exhibit tau pathology in a given region with high confidence. As expected, tau load was found to exhibit a perfect linear relationship with raw SUVR, and was positively related to CSF A*β*42/40 and p-tau217 levels (Fig. 1f,1h, Extended Data Fig.1b). Meanwhile, TPP displayed a saturating trend with these variables (Fig. 1e,1g, Extended Data Fig.1a). TPP showed a strong concordance with Braak staging ^17,19^, whereas SUVR did not (Fig. 1c,1d). Both TPP and SUVR increased monotonically across cumulative Braak stages (Fig. S1). These analyses were repeated using Braak stage regions defined by ^20^, yielding consistent results (Extended Data Fig.1c-j). These findings underscore differences between SUVR and TPP in representing tau pathology, with SUVR indicating quantitative tau load, while TPP reflects a Braak-like distribution pattern. Given that these two measures appear to represent different properties of tau distribution, subsequent analyses are performed across both measures, unless otherwise noted.

**Fig. 1:**
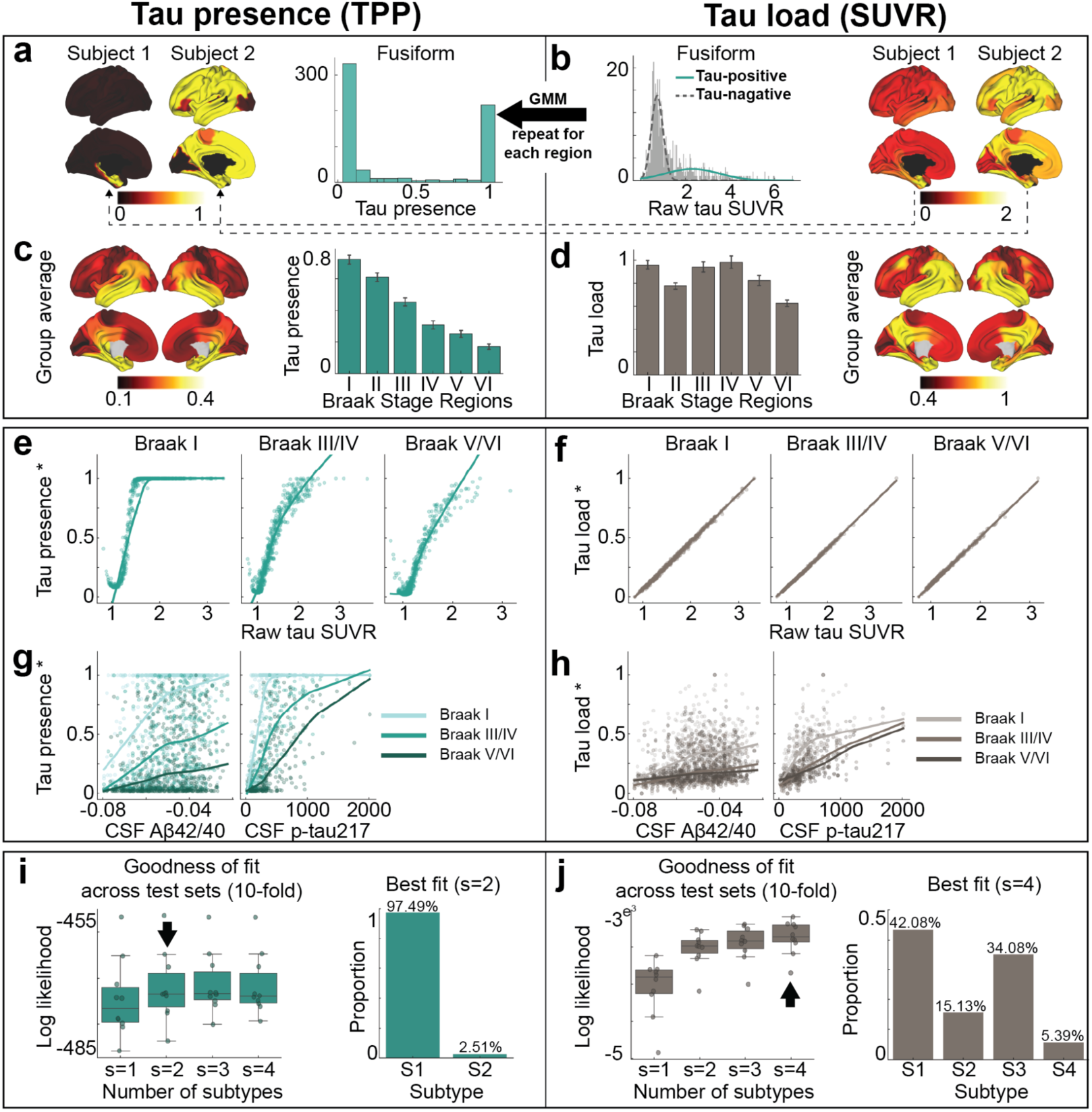
Two distinct tau measurements. **a-d)** Derivation of tau presence (left; where tau is) and tau load (right; how much tau accumulated). **a, b**) Example derivation for each brain region. Histograms depict SUVR and TPP distributions for the left fusiform region. Cortical surface projections illustrate regional tau presence and load for two representative subjects: one with low tau accumulation (Subject 1) and one with high tau accumulation (Subject 2). For each brain region (e.g. left fusiform region): b) normalized tau-PET SUVR represents tau load, scaled relative to 0-1 range defined by cognitively normal A*β*-negative participants; a) tau presence (tau-positive probabilities, TPP) is estimated using a two-component Gaussian mixture model applied to SUVRs across all participants, with TPP defined as the posterior probability of belonging to the higher-SUVR (tauopathy-related PET signal) component. **c, d**) Group-level tau presence and load. Cortical surface projections show regional mean SUVR and TPP values averaged across participants. Bar plots depict mean ± SEM values of SUVR and TPP for Braak stage regions ^19^. **e-h)** Associations between tau presence/load and biological measurements. **e, f)** Relationship with raw SUVR values across Braak stages; **g, h**) relationships with CSF A*β*42/40 and CSF p-tau217 levels. **i, j)** SuStaIn-derived spatiotemporal patterns for tau presence and load. For each subtype model (s=1-4), distributions of average negative log-likelihood across 10 cross-validation folds of left-out individuals are shown, higher values indicate better fit. The best-fit model is marked by the black arrow. Bar plots show the proportion of participants assigned to each subtype. Tau presence yields a single consistent subtype, whereas tau load supports multiple subtypes. SUVR: standardized uptake value ratio; TPP: tau-positive probability; CSF: cerebral spinal fluid; Aβ:beta-amyloid; p-tau: phosporylated tau; SuStaIn: Subtype and Stage Inference.

### Tau load but not distribution differs across individuals

Tau pathology in AD demonstrates substantial variability across the population, with our previous work identifying four distinct patterns of tau accumulation ^28^. We sought to determine whether distinct subtypes exist for both tau presence and tau load, or if these subtypes are observed only in one or the other. To address this, we employed Subtype and Stage Inference (SuStaIn)^91^ models to determine if both tau presence and tau load exhibit distinct subtypes.

Our analysis revealed at least four distinct subtypes for tau load, consistent with previous findings, while tau presence appeared largely consistent across participants (Fig. 1i,1j). Although two subtypes emerged for tau presence, 97.5% of participants were classified into a predominant subtype (S1), with the second subtype (S2) displaying substantial uncertainty, likely influenced by outliers with strong laterality effects (Fig. 1i, S2a, S2b). Importantly, the single-subtype pattern for tau presence was not driven by the soft binarization introduced by mixture modeling. When applying positive-probability measures derived from cognitive data instead, three stable cognitive presence subtypes emerged (Fig. S3). Together, our results suggest that the spatial distribution of tau (tau-positive regions) is highly consistent among individuals, whereas the tau load within those regions shows significant inter-individual variability.

### Classical AD-related elements differently explain tau load and distribution

To investigate hypotheses of tau propagation, we employed the Susceptible-Infectious Recovered (SIR) agent-based model ^77^, a connectome-based disease propagation framework, to simulate tau accumulation across the brain. These models were first fit using group-averaged tau patterns across all Aβ-positive participants (Fig. 2a). The SIR model simulated regional tau synthesis, misfolding, clearance, and spread over a brain connectome, with *a priori* (i.e. measured) regional information influencing model parameters. We first used the SIR model to test whether the observed tau presence or tau load could be explained by propagation along the brain’s anatomical network alone. The model’s performance was evaluated based on its ability to reconstruct the group-averaged regional tau TPP or SUVR across all participants. After testing multiple potential epicenters, the entorhinal cortex emerged as the best-fitting epicenter for tau presence ^17,81,82^, though it was only among the top five candidates for tau load (Extended Data Fig. 2). Based on prior knowledge ^17,72,82^, we selected the entorhinal cortex as the epicenter for subsequent analyses, unless otherwise specified. Our results indicated that diffusion of tau through anatomical connections effectively recapitulated patterns of tau presence (TPP) (R=0.85, MSE=0.02), whereas it did not reproduce tau load (SUVR) as accurately (R=0.45, MSE=0.07) (Fig. 2b,2c). This conclusion was further supported when we applied an alternative connectome-based simulation framework ^72,83^ and a commonly used statistical inference approach ^52,84,85^ (Extended Data Fig. 3).

**Fig. 2:**
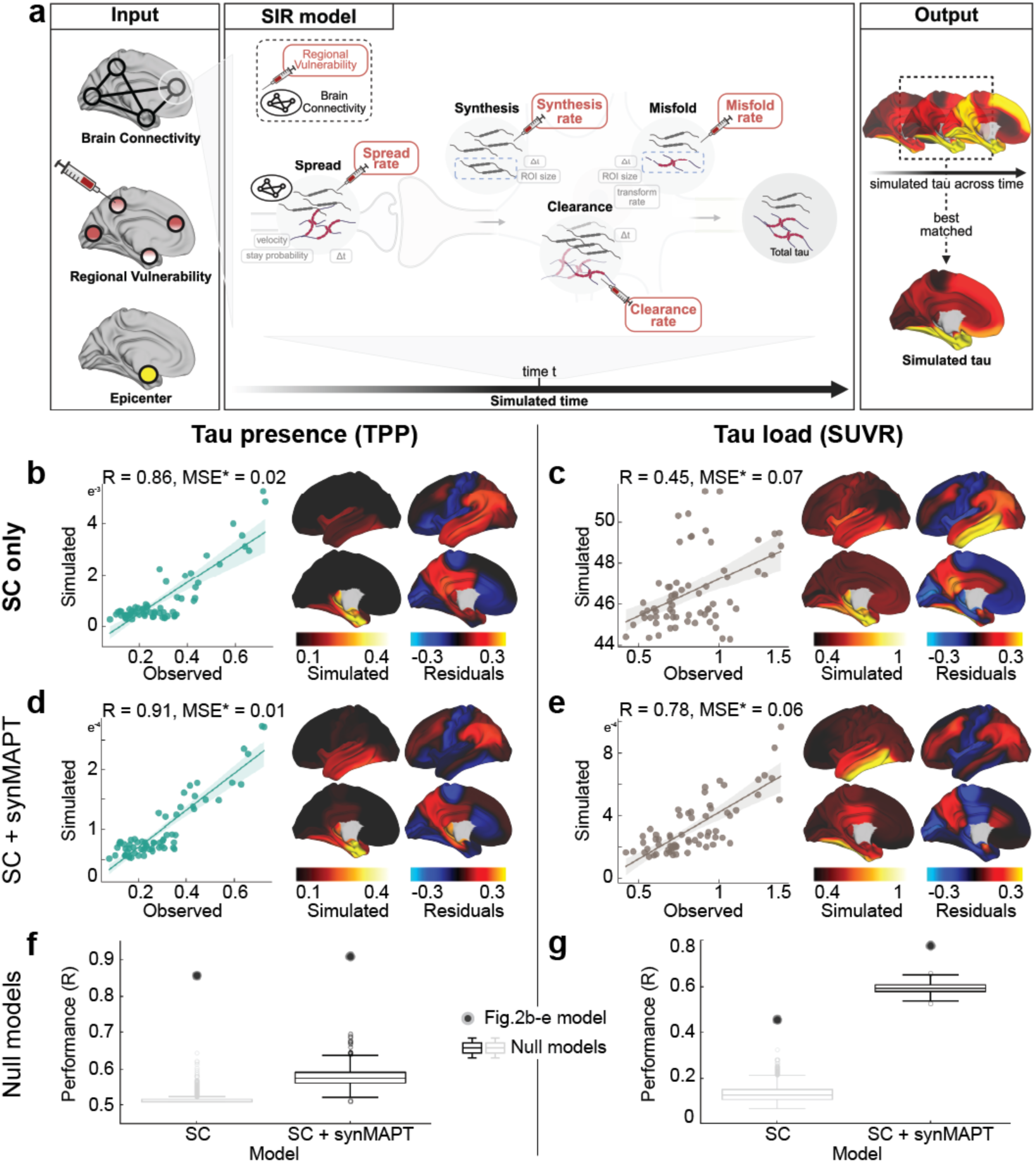
AD-related regional factors improve models of tau load more than models of tau presence. **a)** The Susceptible-Infectious-Recovered (SIR) ^77,78^ model simulates the spread synthesis, clearance, and misfolding of tau. Global parameters affecting each process are shown in grey boxes, while regional biological information is incorporated via key input parameters (red highlights): spread rate, synthesis rate, clearance rate, and misfold rate. **b, c)** SIR model performance for tau presence **b)** and tau load **c)**, simulating tau propagation through a structural connectome derived from healthy young individuals using the entorhinal cortex as the epicenter. Scatter plots compare SIR model simulations to observed regional tau presence/load, averaged over participants, where each dot is a brain region. Cortical surface projections show simulated tau patterns and model residuals. Residuals represent the difference between observed and simulated tau values, with the simulated values normalized to the range of observed values. **d, e)** Influence of regional vulnerability on tau propagation. Three regional properties were incorporated into the model: *MAPT* expression, *APOE* expression, and A*β* deposition. The best-fitting regional property for tau presence (SC plus *MAPT* expression affecting regional synthesis) is shown in **d)**, and for tau load in **e)**. MSE* was calculated using normalized simulated values (scaled to the range of observed tau values) to ensure a fair comparison, as simulated values are not intrinsically bounded. **f, g)** Performance distributions of null models. Box plots show the performance of 1,000 scrambled null models fitted to group-averaged tau presence **f)** and tau load **g)**. Null models included (i) scrambled structural connectivity (SC) only and (ii) scrambled SC combined with regional MAPT expression used as the synthesis rate (SC+synMAPT). Structural connectivity was scrambled while preserving weight and degree distributions. SUVR: standardized uptake value ratio; TPP: tau positive-probability; SC: structural connectivity; R: Pearson correlation coefficient R; MSE: mean squared error; *MAPT*: microtubule associated protein tau; *APOE*: Apolipoprotein E; syn: synthesis rate; clear: clearance rate; spread: spread rate.

Next, we examined the influence of three well-established AD-related factors on tau propagation - A*β* deposition level from [18F]-flutemetamol-PET, Apolipoprotein E (*APOE*) and microtubule-associated protein tau (*MAPT*) gene expression from the Allen Human Brain Atlas (AHBA) ^86^ - on regional susceptibility to both tau presence and load. This was done by allowing each factor to have a regionally-varying effect on either tau synthesis, clearance, spreading, or misfolding rates of tau propagation, based on levels of the factor in a given region. For each regional factor, the best-performing rate was selected and considered as the mechanism through which the factor most strongly influences tau propagation (Supplementary Table S2). Among these models, allowing regional *MAPT* expression to influence tau synthesis rate (presence ΔR=0.0523, load ΔR=0.3243) or regional A*β* to influence tau spread rate (presence ΔR = -0.1727, load ΔR = 0.1150) improved predictions of tau load more than tau presence (Fig. 2d-2g). Regional *APOE* expression did not enhance predictions for either tau presence or tau load. Similar results were observed when fitting the model with a different anatomical connectome derived from a separate sample of healthy young individuals ^87,88^ (62 cortical regions, excluding the hippocampus and amygdala (Fig. S4, Supplementary Table S3).

When anatomical connectivity was disrupted, model performance for tau presence deteriorated substantially and was only minimally recovered by incorporating regional MAPT expression (Fig. 2f). In contrast, even under disrupted connectivity, predictions of tau load substantially improved when regional MAPT expression was included (Fig. 2g). This underscores the importance of brain connectivity in explaining tau presence in our model, and local factors being more important in explaining tau load.

Together, these results suggest that Braak-like tau presence, especially in Braak I-IV regions, is primarily driven by diffusion along anatomical connections, while regional tau load is best explained by a combination of anatomical connectivity and regional *MAPT* expression or A*β* deposition.

### Connectome alone shapes tau presence, while regional vulnerability influences tau load

Tau is thought to propagate from cell to cell through synaptic connections and is likely influenced by conventional AD-related properties, like Aβ. However, other properties might determine where and how much tau accumulates in one region and other hypotheses have not been ruled out. Rather than limiting our focus to a small number of predefined factors of interest, we performed an exploratory analysis across eight types of normative connectomes ^87^ (Fig. 3a) and 49 regional biological factors ^89–94^ (Extended Data Fig. 4), assessing their contributions to tau propagation using SIR models and testing multiple hypotheses of tau propagation. These included various measures of brain connectivity, encompassing both anatomical connections derived from probabilistic tractography of DWI data, but also inter-regional similarity, which reflects potential interactions between regions based on shared molecular, cellular, and functional properties. Inter-regional similarity, derived (separately) from gene expression, neurotransmission marker density, cellular morphology, glucose metabolism, haemodynamic activity, and electrophysiology, captures patterns where regions with similar neurobiological profiles may be similarly affected by tau pathology. The analysis also included diverse regional intrinsic properties such as morphometric qualities, neurotransmission marker profiles, metabolic measures, cell-type distributions, and AD-related genetic profiles. Overall, we tested 1,576 models, consisting of eight connectome-only models and 1,568 models combining connectomes with regional biological factors (8 connectome *×* 49 regional factors *×* 4 tau mechanistic rates). For visualization, we selected the eight connectome-only models and 392 connectome-regional factor models, where the latter combined 8 connectomes with the 49 regional factors, each paired with the best-performing tau mechanistic rate (i.e., synthesis, clearance, misfold, spread).

**Fig. 3:**
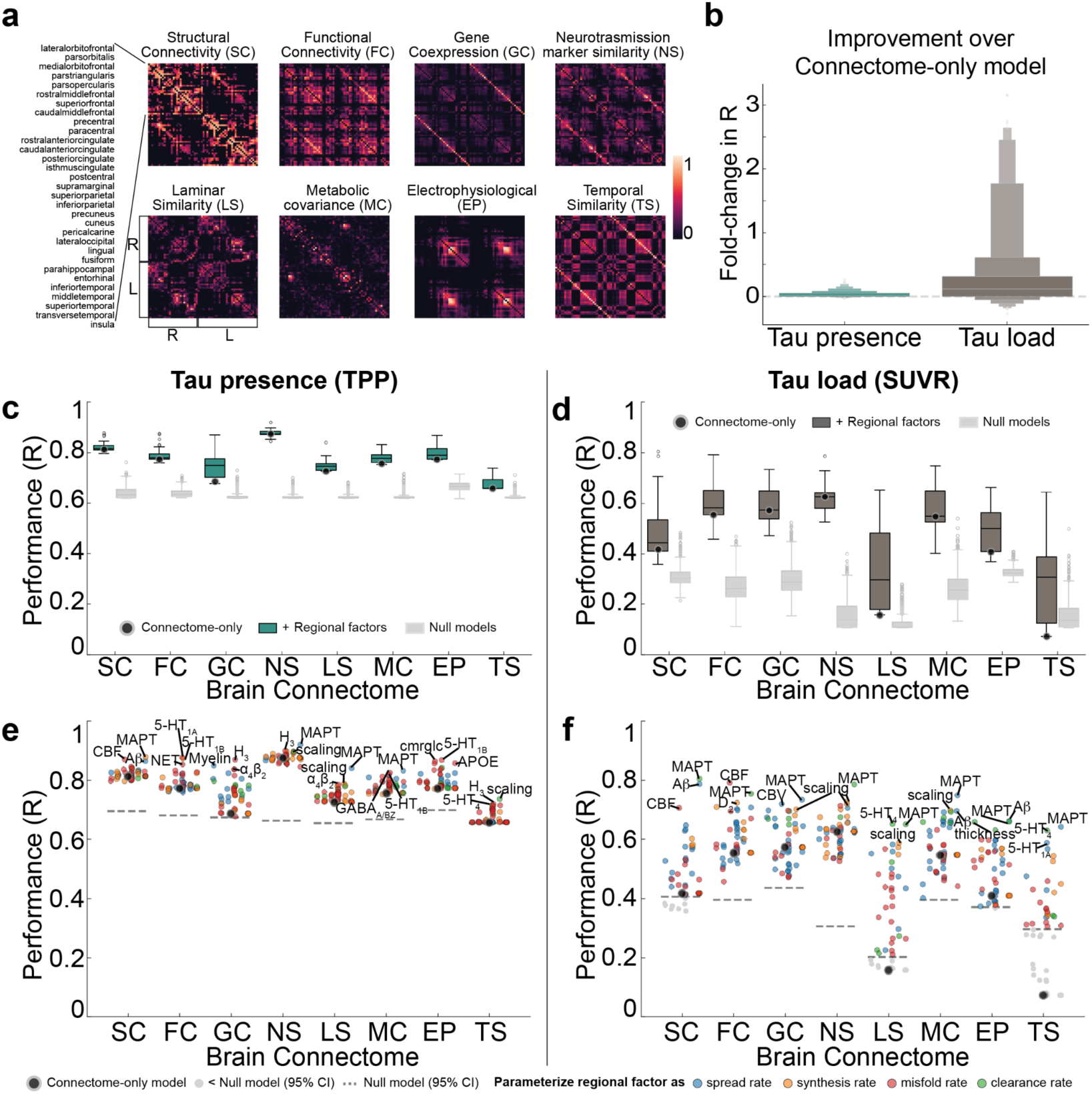
Distinct connectome and regional properties shape tau presence and load. **a)** The SIR model was fit using eight different types of normative connectomes representing different types of regional relationships in the brain, extracted from ^87^. **b)** Across 400 selected models, tau load (SUVR) models were more improved by the addition of regional factors than tau presence (TPP) models were. The boxenplot shows the distribution of fold-change increase in model performance across all models. **c, d)** Performance of models across different types of connectomes fitted to c) TPP and d) SUVR data. Across all connectomes, most connectome-only models (indicated by larger black circles) exceeded chance (95% CI of 1000 models scrambling connectomes but preserving weight and degree distribution). **e, f)** Performance of individual models fitted to tau TPP e) and SUVR f), respectively. Each dot represents one model. The dot color indicates what rate parameter the regional information was influencing. Grayed-out dots did not exceed null model performance (indicated by dashed lines). The regional factor added to the model is labeled for the top 3 models of each connectome. SUVR: standardized uptake value ratio; TPP: tau positive-probability; R: Pearson correlation coefficient R; SC: structural connectivity; FC: functional connectivity; GC: gene coexpression; NS: neurotransmission marker similarity; LS: laminar similarity; MC: metabolic covariance; EP: electrophysiological; TS: temporal similarity; CBF: cerebral blood flow; CBV: cerebral blood volume; thickness: cortical thickness; scaling: allometric scaling.

We first investigated the influence of regional vulnerability on tau presence and tau load by calculating the fold-change improvement in model performance when regional biological factors were incorporated compared to the connectome-only model. Overall, tau load (SUVR) models (mean=0.36-fold improvement, std=0.68, range=[-0.27, 3.15]) benefited more from the inclusion of regional factors compared to tau presence (TPP) models (mean=0.03, std=0.05, range=[-0.02, 0.27]) (Fig. 3b). Model performance was evaluated against null distributions generated by simulating models through scrambled connectomes. We found performance of connectome-only models to exceed chance levels for all tau presence models, and all tau load models except those using laminar similarity and temporal similarity networks. When looking at general tendencies across models, tau presence was best recapitulated by diffusion through neurotransmission marker similarity and anatomical networks on average (Fig. 3c), while tau load was best explained by diffusion through neurotransmission marker similarity and functional connectivity networks (Fig. 3d). Our findings confirm that anatomical connectivity is a key factor in modeling tau presence, though the finding that a neurotransmission marker similarity network provided the best fit for modeling both tau presence and load was unexpected, and is further explored below.

Models involving spread through anatomical connections, with *MAPT* expression influencing regional tau synthesis or misfolding, A*β* influencing regional tau spread, or cerebral blood flow influencing regional tau clearance, were among the top-performing models explaining both tau presence and tau load. These findings align with widely hypothesized mechanisms, supporting the validity of our model. Moreover, several less-explored tissue properties emerged as important for moderating tau load and distribution. For tau presence, the top 10 best-fitting models predominantly involved diffusion through a neurotransmission marker similarity network (Table 1). Notable tissue properties included allometric scaling influencing tau synthesis or misfolding rates, as well as different neurotransmission markers modulating tau clearance or synthesis (Fig. 3e, Extended Data Fig. 5a left). In contrast, the best-fitting models for tau load featured diverse combinations of brain connectivity and regional factors (Table 1). Specifically, various forms of brain connectivity coupled with regional *MAPT* expression affecting tau misfolding or spreading provided the most accurate fits (Fig. 3f, Extended Data Fig. 5a right). Across all models and irrespective of the chosen brain connectome, the regional biological factors that improved models of tau presence most on average were serotonin 5-HT_1B_ receptor, *MAPT*, and allometric scaling, whereas for tau load, the key factors were nicotinic acetylcholine receptor (*α*_4_β_2_), *MAPT*, and serotonin 5-HT_4_ receptor (Extended Data Fig. 5b,c, Supplementary Table S4). Overall, these findings reaffirm that the spatial pattern of tau propagation is governed primarily by a connectome-based mechanism, whereas tau accumulation within affected regions is modulated by both network connectivity and intrinsic regional properties.

**Table 1:**
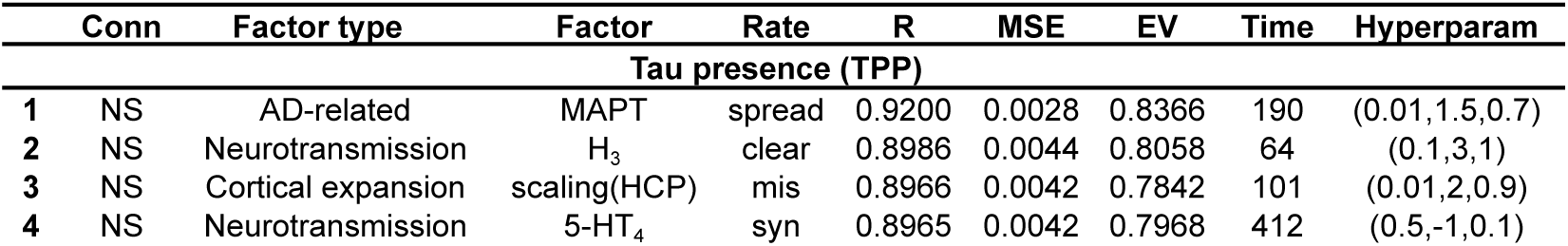
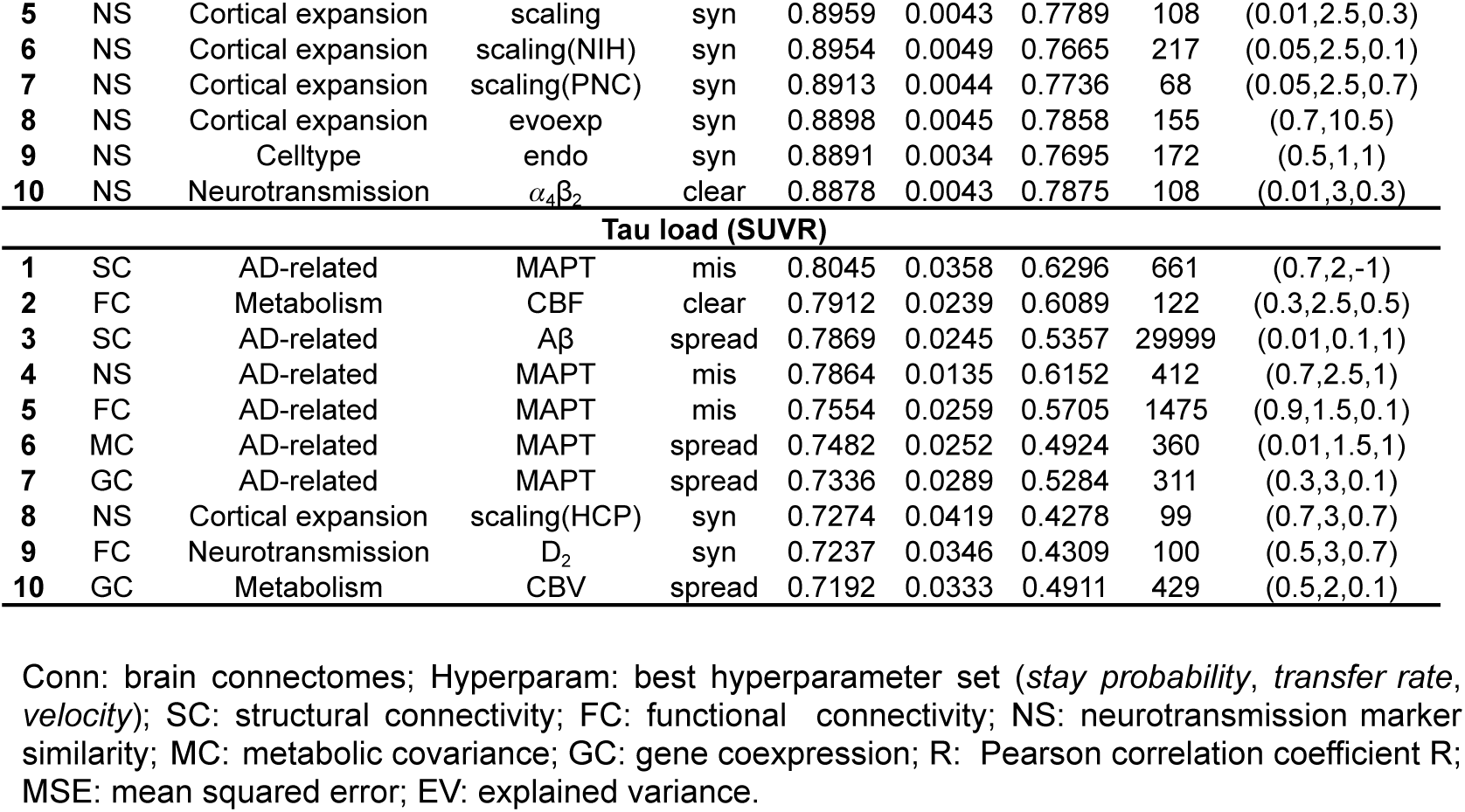
Top 10 models for tau load and presence.

### Distinct associations of different neurotransmission markers in tau patterning

Our results indicated that neurotransmission marker distribution (i.e., the neurotransmission marker similarity network, see Methods) might play a previously underappreciated role in explaining tau propagation. To delve deeper, we aimed to identify the specific neurotransmission marker profile most associated with tau accumulation patterns.

We first investigated the relationship between the neurotransmission marker similarity network and the structural connectivity network. The neurotransmission marker network closely resembled the anatomical connections derived from DWI, but with notable differences (Fig. 4a). Specifically, the neurotransmission marker similarity network displayed stronger contralateral correlations, whereas the anatomical network was dominated by stronger ipsilateral connections. Compared to anatomical connections, the neurotransmission marker similarity network exhibited stronger connections in the inferior parietal, medial orbital frontal, superior temporal, and precuneus regions, while showing weaker connections in the inferior temporal, insular, and entorhinal regions.

**Fig. 4:**
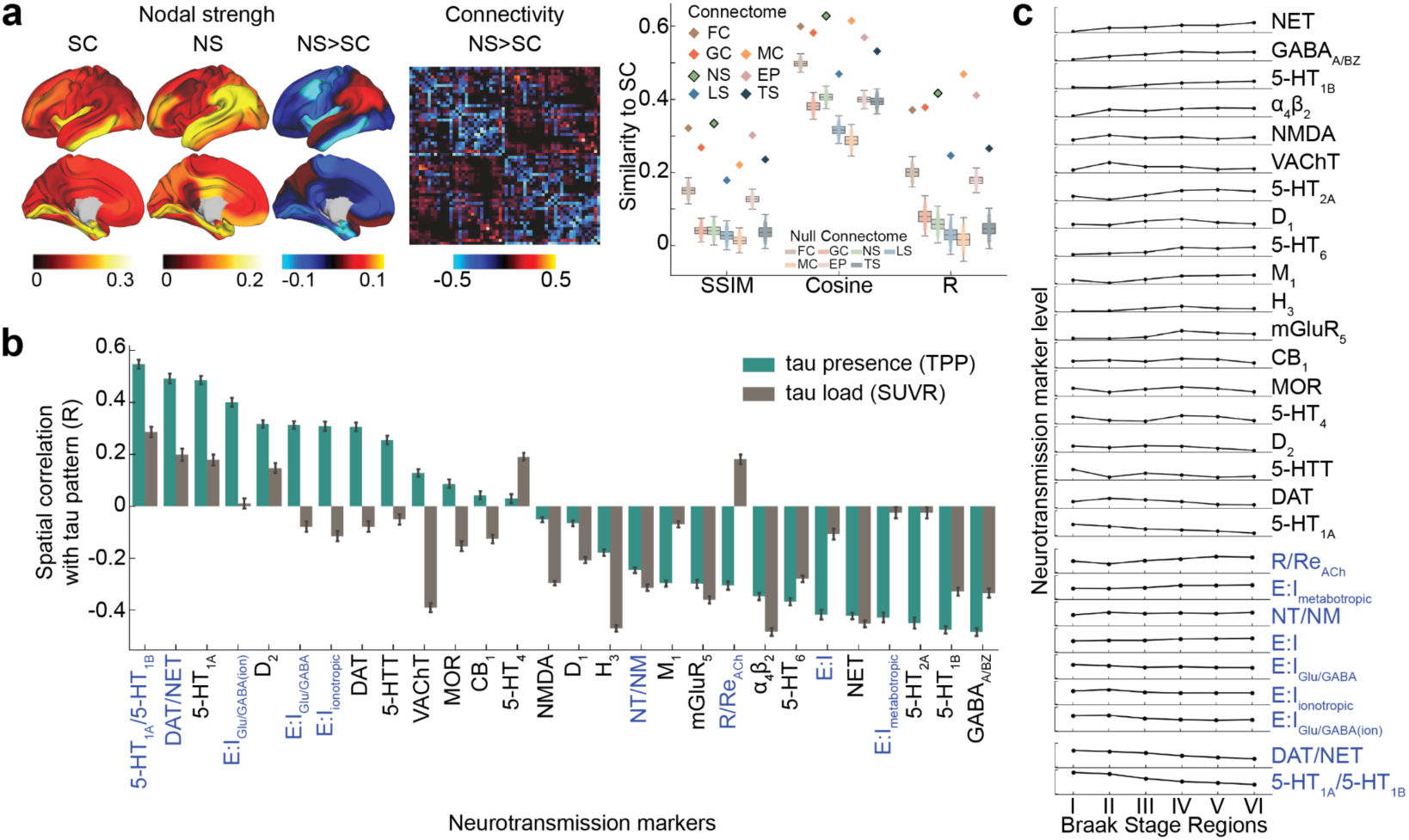
Neurotransmission marker similarity mirrors structural connectivity, while neurotransmission marker distribution aligns with tau patterns. **a)** Cortical surface projections show nodal strength for structural connectivity, neurotransmission marker similarity, and their differences. A heatmap illustrates the connectivity differences between neurotransmission marker similarity and structural connectivity. The right panel compares the similarity (assessed using three metrics: SSIM, cosine similarity, and Pearson correlation coefficient R) between structural connectivity and seven other tested brain connectomes (diamond). The similarity between structural connectivity and neurotransmission marker similarity is highlighted with a black box. Similarity between structural connectivity and null connectomes (1,000 connectivity matrices per connectome, scrambled while preserving weight and degree distributions) is shown with light boxplots. **b)** Spatial correlation of neurotransmission markers (black text) and derived ratios (blue text) distributions with tau presence (TPP, green) and tau load (TPP, grey) across 66 brain regions used in previous analyses ^80^. **c)** Neurotransmission marker distributions across regions grouped by Braak stages, with neurotransmission markers and derived ratios sorted separately by the magnitude of change from regions assigned to Braak stage I to those assigned to stage VI. NS: neurotransmission marker similarity; SC: structural connectivity; FC: functional connectivity; GC: gene coexpression; LS: laminar similarity; MC: metabolic covariance; EP: electrophysiological; TS: temporal similarity; SSIM: structural similarity index; Cosine: cosine similarity; R: Pearson correlation coefficient R; TPP: tau-positive probability; SUVR: standard uptake value ratio; E:I: excitatory-to-inhibitory ratio; Glu: glutamate receptors; GABA: gamma-aminobutyric acid receptors; ion: ionotropic; NT/NM: neurotransmitter-to-neuromodulator ratio; R/ReACh: acetylcholine receptor-to-reuptake ratio.

To better understand the pattern of neurotransmission markers that explain the efficacy of neurotransmission marker similarity networks in propagating tau in our models, we investigated which specific neurotransmission marker distribution most closely resemble observed tau patterns. We investigated neurotransmission marker distributions and their combinations, derived both *a priori* from previous literature ^95–98^ and *a posteriori* based on observations from this study. Our findings showed that tau presence and Braak staging were highly correlated with the ratio between serotonin receptors 1A and 1B (5-HT_1A_/5-HT_1B_), dopamine and norepinephrine transporter densities (DAT/NET), receptor to reuptake ratio of acetylcholine (R/Re_ACh_), and the ratio of excitatory to inhibitory (E:I) receptors (Fig.4b, Fig.4c). Interestingly, the E:I ratio exhibited complex relationships: ionotropic (E:I_ionotropic,_ E:I_Glu/GABA:ion_) and glutamate to GABA (E:I_Glu/GABA_) receptor ratios correlated positively with tau presence and Braak stage, whereas the metabotropic E:I receptor ratio (E:I_metabotropic_) showed a negative correlation. Conversely, tau load was most strongly associated with the distribution of histamine receptors (H_3_), nicotinic acetylcholine receptors (*α*_4_β_2_), the norepinephrine transporters (NET) and vesicular acetylcholine transporter (VAChT) (Fig. 4b).

### Different factors explaining tau subtypes

As previously noted and demonstrated above, variation in the pattern of tau accumulation has been described across the population. We aimed to explore whether biological or connectome-based factors contribute to these differences. Building on our previous work ^28^, we extracted the mean regional tau load (SUVR) for each tau-PET subtype (Fig. 5a) and employed SIR models to explore factors contributing to tau load pattern across the four subtypes, utilizing the same eight brain connectomes and 49 biological factors from the prior analysis.

**Fig. 5:**
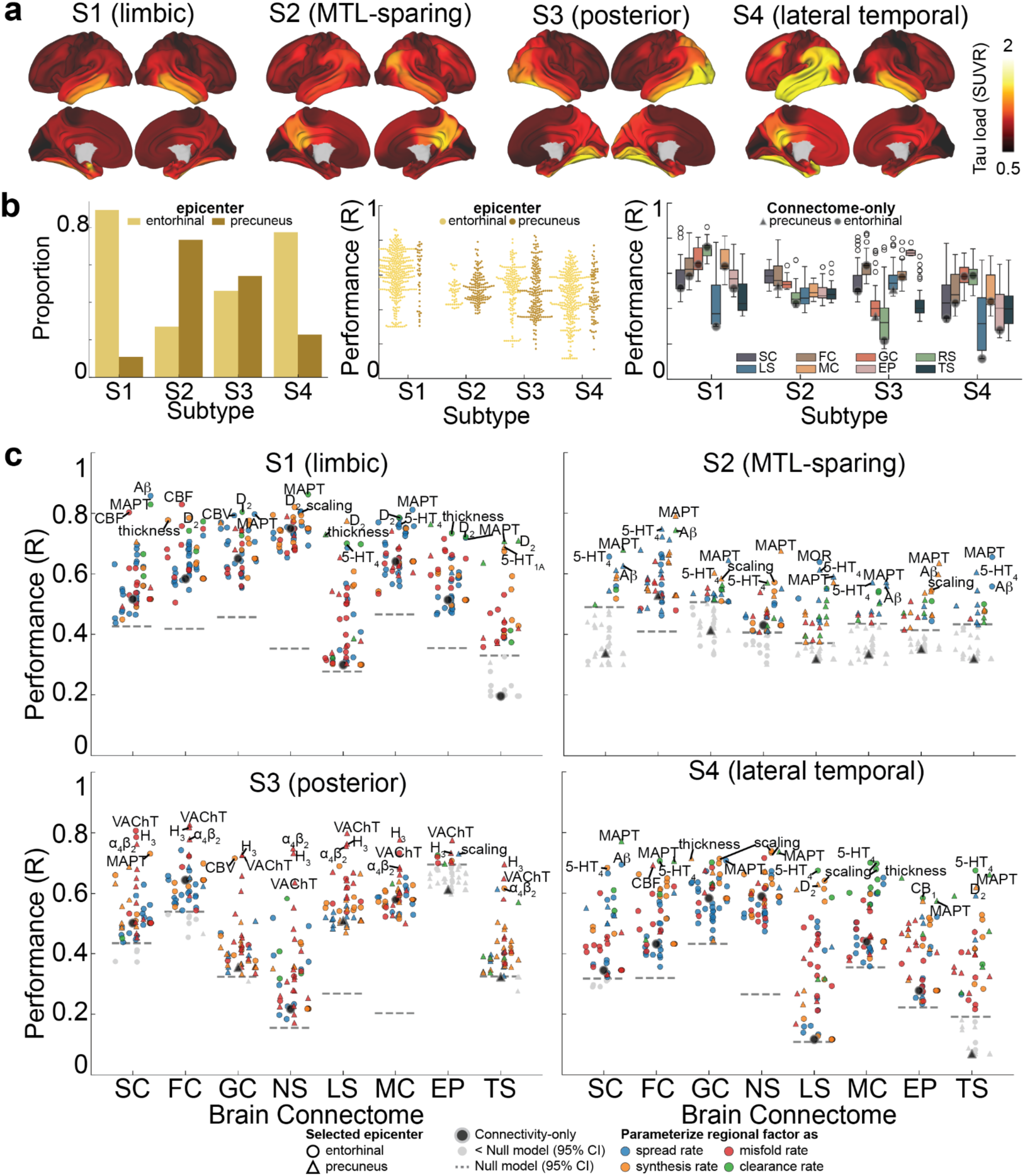
Tau load varies across individuals, driven by distinct factors. **a)** Average tau SUVR pattern for each subtype, extracted based on our previous work ^28^. **b, c)** The SIR model was fit for each of the four SUVR subtypes using eight different normative connectomes and 49 different regional biological factors, as in Fig. 3a, Extended Data Fig. 4. Only models better than chance were shown (95% CI of 1,000 models scrambling connectomes but preserving weight and degree distribution). **b)** The proportion (left) and performance (middle) of models with best-fitting epicenters in entorhinal yellow) and precuneus (brown) regions; and (right) performance of models across different connectomes. **c)** Performance of individual models fitted to each of the four SUVR subtypes. Each dot represents one model, with color indicating what parameter the regional information was influencing. Grayed-out dots did not exceed null model performance (95% CI, indicated by dashed lines). The regional factor added to the model is labeled for the top 3 models of each connectome. SUVR: standardized uptake value ratio; TPP: tau positive probability; SC: structural connectivity; FC: functional connectivity; GC: gene coexpression; NS: neurotransmission marker similarity; LS: laminar similarity; MC: metabolic covariance; EP: electrophysiological; TS: temporal similarity, R: Pearson Correlation coefficient R; CBF: cerebral blood flow; CBV: cerebral blood volume; thickness: cortical thickness; scaling: allometric scaling.

Building on earlier findings indicating distinct epicenters across individuals ^28^, we tested the model by varying the epicenter between the entorhinal cortex and the precuneus for each of the four subtypes. The entorhinal cortex emerged as the best fitting epicenter for most models of the limbic and lateral temporal subtypes, while the precuneus was the best-fitting epicenter for most models of the MTL-sparing subtype. For the posterior subtype, both the precuneus and entorhinal cortex were identified at similar rates (Fig. 5b, left). Notably, tau patterns in all subtypes besides the MTL-sparing subtype could be well captured (i.e. among top 10 models) using the entorhinal cortex as the epicenter and diffusion along anatomical or functional connections (Fig. 5b middle and right, Supplementary Table S5). For the MTL-sparing subtype, however, the models consistently identified the precuneus as the best-fitting epicenter, with tau propagation more accurately modeled through brain functional connectivity (Fig. 5b right, Supplementary Table S5).

To further elucidate the factors shaping tau load across subtypes, we identified and summarized the top-contributing factors (Fig. 5c, Supplementary Table S5, S6). Consistent with established disease mechanisms, tau diffusion along anatomical connections, and modulated by regional *MAPT* or A*β* expression, effectively explained all subtypes (Fig. 5c). The limbic subtype (S1) exhibited contributory factors that closely resembled those influencing general tau load, primarily driven by propagation through neurotransmission marker similarity networks or anatomical connections, and regional intrinsic properties linked to conventional AD-related factors (Fig. 5c top left). Notably, the distribution of dopamine receptors (D_2_) played a potential role in tau accumulation in this subtype. For the MTL-sparing subtype (S2), tau load appeared to originate from the precuneus, spreading through brain functional connections and influenced predominantly by conventional AD-related factors and distribution of serotonin (5HT_4_), as well as the extent of allometric expansion (Fig. 5c top right). Interestingly, looking at connectome-only models, only the functional and neurotransmission marker similarity models exceeded chance levels of performance. In the posterior subtype (S3), tau load was consistently influenced by the regional distribution of three neurotransmitter receptors, H_3_, VAChT, and *α*_4_β_2_, primarily affecting tau clearance (Fig. 5c bottom left). This underscores the potential role of acetylcholine and histamine signaling in modulating tau load in the posterior subtype. Lastly, the lateral temporal subtype (S4) exhibited a mechanism similar to that of the limbic subtype (S1), though a distinct pattern of models did not clearly emerge for this subtype (Fig. 5c bottom right). Extended Data Fig. 6 shows the regional biological factors that contributed most across all models, and independent of the connectome used, for each subtype.

## Discussion

Our work suggests tau presence and tau load represent two distinct phenomena mediated by different underlying mechanisms (Fig. 6). Tau presence, i.e., where tau accumulates in the brain, follows a Braak-like pattern, primarily driven by diffusion along anatomical connections, and this distribution pattern is consistent across individuals. In contrast, tau load, i.e., how much tau accumulates in a given region, deviates from the Braak pattern and depends on a combination of brain connectivity and regional biological properties, with substantial variability observed between individuals. In this study, we approximated tau presence through the reach of tau pathology into a region, measured using a tau positive probability (TPP). We equate this approach to tau seeding activity, indicative of the initial presence of misfolded tau capable of inducing further aggregation. Tau seeding activity has notably been shown to precede the appearance of overt tau pathology, to serve as a critical driver of its propagation ^99,100^, and to follow synaptic transmission pathways ^99,101,102^, consistent with the idea that tau presence primarily spreads along neuronal networks. In contrast, tau load appears moreso to be influenced by local biological factors, especially *MAPT* expression levels and A*β* deposition. These findings together support a model where tau seeds propagate through anatomical connections in a stereotyped fashion early in the disease process ^103^, whereafter tau misfolding accelerates in A*β*-prone regions with abundant *MAPT*, and is further modified by person-specific brain properties. This model provides a mechanistic account supporting previous models of A*β*-induced tau spreading and individual variability in tau accumulation. More importantly, these findings suggest a duality in tau spread mechanisms, where the spatial pattern of tau propagation is governed by different mechanisms from those that dictate local tau accumulation within affected regions.

**Fig. 6:**
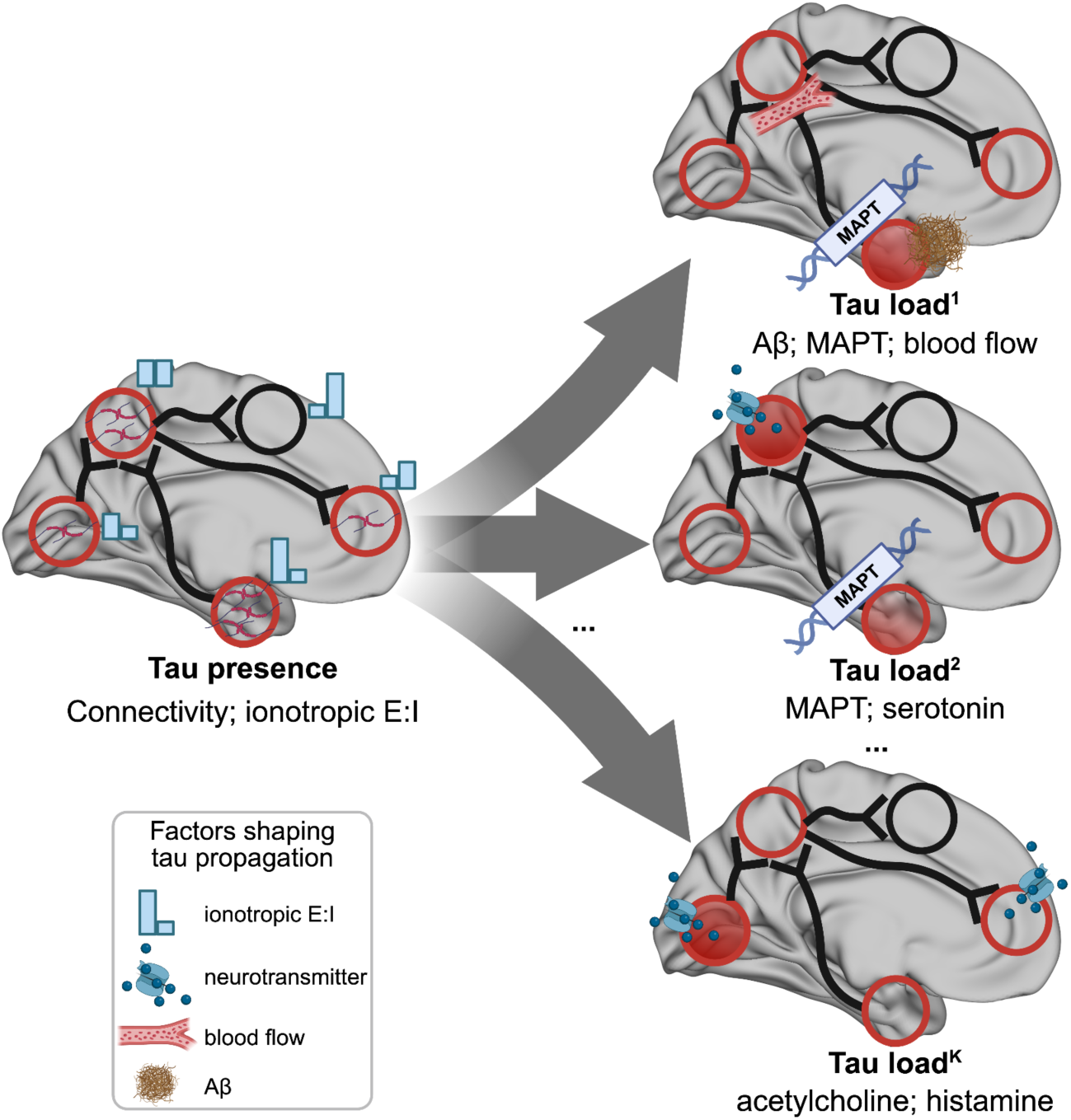
A theoretical model summarizing the spread of tau pathology in AD. The model delineates two distinct tau spread phenomena mediated by different underlying mechanisms: tau presence (where tau appears) and tau load (how much tau accumulated). Tau presence emerges consistently across individuals and is primarily driven by diffusion along anatomical connections, and modulated by ionotropic excitation-inhibition (E:I) balance. In contrast, tau load exhibits substantial inter-individual variability and reflects the combined influence of brain connectivity and multiple regional biological factors, encompassing distinct vulnerability mechanisms.

The distinction between these two properties of tau accumulation, presence and load, holds implications for both the quantification of tau and investigation into the cellular vulnerability to tau. Braak staging, the classical method of measuring the progression of tau pathology postmortem, assigns stages based on whether tau pathology has progressed to a given region ^18,61^, making it a measurement of tau presence rather than tau load. Instead, tau load is traditionally quantified postmortem using semi-quantitative scales (i.e. sparse, light, moderate, severe) that are not necessarily standardized across neuropathologists ^104–106^. In contrast, most tau-PET studies use SUVR as a fairly standard continuous measure of tau load ^5,107,108^, including in clinical trials ^109–111^. Recent end-of-life comparison studies suggest that tau-PET scans do not become “positive” until postmortem Braak stage IV-V (of VI) ^112–114^, though this may highlight a scenario where the distinction between tau presence and load is important. Braak stage IV may indicate that there are one or more tau inclusions present in stage IV regions, but does not necessarily indicate how much tau is in the brain. Supporting this notion, in the present study, our measure of tau presence showed a Braak-like distribution while tau load (SUVR) did not. In addition, greater agreement between in vivo and postmortem measures of tau has been found when both use measures of tau load ^115,116^. These findings highlight that presence and load are distinct properties of tau accumulation that can both be quantified in vivo using tau-PET. This notion has recently become more appreciated ^117–119^, though tau presence in these studies is often termed as spatial spread, extent or progression. While not the main focus of the present study, other recent studies have done well to indicate how tau presence and load may be displaced in time and in relation to other disease markers ^118–121^, and may therefore be of great interest as primary or secondary outcomes in upcoming drug trials for tau-targeting therapies.

The implications of our study findings suggest that tau presence and load are representative of distinct aspects of cellular or regional vulnerability, and may be mediated by differing biological properties. Specifically, whether cells in a population are susceptible to accumulating pathological tau, and how much tau accumulates in that population, may be dissociable phenomena. In the present study, tau presence was strongly related to connectivity to the entorhinal cortex, whereas tau load was best explained by a combination of connectivity and other tissue properties. We did not find appreciable population variation in regional tau presence, despite obvious variation in regional tau load. This is consistent with neuropathology studies, which have noted that, e.g., “hippocampal-sparing” cases do show tau pathology in the hippocampus, just less than would be expected given global tau load ^122,123^. Based on these data, neurons susceptible to the accumulation of AD tau are likely consistent across individuals, and alongside a wealth of literature on the topic ^56,58^, we speculate this susceptibility is driven by cell-intrinsic factors. Supporting this idea, neurons in animals that do not natively accumulate AD tau pathology will still accumulate AD tau when exposed to AD-like conditions, such as A*β* deposition *and MAPT* expression ^38–40,124,125^. This points to the idea that certain neurons are capable of expressing pathological tau, and will do so under pathological conditions. Our simulation model supports this, by showing that connectivity-driven spread of AD tau with prion-like features into regions with tau-susceptible neurons can recapitulate the pattern of tau presence across the population. However, this simulation was not sufficient to explain population patterns of regional tau load, which we speculate is driven by a combination of cell-intrinsic and cell-extrinsic phenomena that are person specific (discussed further below). The distinction is potentially informative toward the scale of investigation future studies of tau should engage in. Single-cell studies ^126–128^ may be adept at further studying the neuronal susceptibility to tau that underlies tau presence, whereas spatial omics and neuronal circuit studies may better capture the vulnerable features of cell populations that may drive tau load ^129,130^.

The present study consistently showed that the spread of tau from the entorhinal cortex through anatomical connections, and moderated by regional *MAPT* or A*β*, best explained tau accumulation patterns. These findings, especially juxtaposed against the large number of exploratory simulations included in this study, strongly support the conventional model of AD. A*β* has long been hypothesized as the key mediator of the fulminant tauopathy characteristic to AD ^4,125,131,132^. Previous *in vivo* human studies have leveraged connectome-based models to support this notion, showing that adding regional A*β* information improves models of tau spread ^51,72,79^. However, the mechanism by which A*β* potentiates tau pathology is still unclear. Our simulations showed that allowing regional A*β* to moderate spread of seed-competent tau, as opposed to production or misfolding of tau, best explained tau patterns across the AD population. These models therefore suggest that the presence of A*β* in a brain region increases the likelihood and/or velocity that tau spreads to that region. This is supported by *in vivo* studies in both animals ^133^ and indirectly in humans^52^, the latter of which suggests distal A*β* stimulates activity-dependent release of tau into A*β*-containing brain areas. Simulations incorporating regional *MAPT* gene expression were also consistently high-performing. Previous studies have noted the correspondence between *MAPT* gene expression and *in vivo* tau deposition patterns ^59,134,135^, and have noted that it adds complementary information to connectome-based models of tau spread ^76,79^, though not in mice ^136^. The present study uniquely incorporates *MAPT* gene expression into a mechanistic simulation of tau spread. The best results came from models where regions with more *MAPT* transcripts misfolded tau at a higher rate. Models allowing *MAPT* to moderate tau synthesis rate (instead of misfold rate) also performed well but were slightly worse. This is interesting given that many *MAPT* mutations increase tau aggregation ^137,138^, perhaps implicating this mechanism as most critical for driving tauopathy. Notably, models incorporating regional *MAPT* or A*β* both performed well independently of one another. Tau propagation models including A*β* were sufficient to explain tau presence patterns, but were not necessary. This may have implications for therapeutic development, which may benefit from targeting both A*β*-lowering and tau misfolding mechanisms.

While the exhaustive exploratory simulations conducted in the present study largely support conventional hypotheses of tau spread, there were several other high performing models that emerged. These particular simulations may nominate additional mechanisms for further consideration as independent or interacting features of AD pathogenesis. One model that performed almost as well as the conventional AD models involved increased tau clearance in regions with greater regional cerebral blood flow (CBF). The glymphatic system is known to be critical for the clearance of AD pathology ^139–141^. Conversely, later stages of AD are known to involve decreased cerebral blood flow ^142–145^, and neurovascular disease reduces resilience to AD pathology ^146–149^. The high performance of this model is particularly interesting given that regions with greater cerebral blood flow largely overlap with regions that experience greater A*β* aggregation, but the effects on tau are in opposite directions. This may point to a competitive complex of A*β*-driven spread and glymphatic clearance of tau in AD-prone regions, the latter of which is likely overwhelmed in later disease stages. These results point to a continual development of glymphatic-aiding therapies, perhaps in addition to A*β*- and tau-targeting drugs.

Simulations incorporating allometric scaling of brain surface area were also surprisingly frequent among top models. Most of these models involved a faster rate of tau misfolding in regions that expand disproportionately with increased brain size. Such regions are under increased metabolic strain and strongly relate to patterns of phylogenic and ontogenic expansion of the brain ^150^. These regions also overlap considerably with regions prone to A*β* aggregation, which may be a less compelling explanation of such models. However, some evidence has supported a “last-in first-out” hypothesis where late-maturing or phylogenetically new areas are more susceptible to aging and AD ^63,151–153^, and our models may provide some hints in support of this idea.

Equally notable were some biological properties previously associated with tau that did not enhance models of tau spread in our study. Cortical myelination has long been associated with decreased tau pathology, stemming from the observation that tau-prone regions in the entorhinal cortex are thinly myelinated while heavily myelinated regions like the primary somatosensory and motor areas are resistant to tau pathology ^153–155^. Recent neuroimaging studies have supported this idea ^62,156,157^. However, simulations incorporating regional myelination were mostly absent from top models in our study. One possible explanation is that cortical myelination patterns are highly related to the brain’s connective topology, rendering myelination redundant in simulations of tau spread over brain networks. Alternatively, cortical myelination patterns also bear strong spatial similarity to a number of other relevant brain properties ^158^, including A*β* deposition patterns, glucose metabolism and cortical expansion. These spatial associations could create incidental associations with tau, but which do not necessarily bear out as relevant in fully simulated mechanistic network models from the present study. Another surprising finding running counter to previous studies ^76^ was that we did not find expression of AD-related genes (other than *MAPT*) or cell-type distributions to improve models of tau spread. Notably, *MAPT* expression, which has a strong a priori association with tau, was sourced from the same dataset, suggesting this null finding was not a methodological issue. This lack of findings suggests that genes regulating intrinsic vulnerability to tau are not necessarily the same as genes affecting AD risk. It is also possible that there are some spatially-dependent effects on tau, but they cannot be detected at our current scale of investigation.

We investigated tau spread over various networks, encompassing both anatomical connections and inter-regional similarity. This approach allows us to test different hypotheses of tau spread and different views of the brain’s communication network beyond direct anatomical connections. Previous studies have consistently found that spread over anatomical networks better explains tau than spatial diffusion ^70–72^, and often ^43,44,159^, but not always ^46,76,160^, spread through functional networks. In the present study, two of the top three models for tau load did indeed use anatomical connectivity as the underlying connectome. However, all of the top-performing models for tau presence surprisingly involved a neurotransmission marker similarity network as the underlying connectome, where regions with neurotransmission marker profiles more similar to the entorhinal cortex were more likely to express tau sooner. One possible explanation is that the neurotransmission marker similarity and anatomical connectivity networks are highly similar, but that the neurotransmission marker similarity network may contain additional molecular information relevant to tau vulnerability. Indeed, previous studies have shown that neurotransmission marker similarity strongly correlates with both diffusion-derived structural connectivity and fMRI-derived functional connectivity ^90^. At our scale of investigation, regional anatomical connectivity is summarized across various tracks, perhaps losing some information. Brain regions with similar neurotransmission marker profiles are likely also communicating (suggesting brain connectivity), and the neurotransmission marker similarity network may therefore act as a functionally-enriched anatomical connectivity network. There is an alternative possibility, where the neurotransmission marker similarity network may reflect differential molecular vulnerability to tau accumulation, and spread of tau represents spontaneous pathological events occurring over a gradient of vulnerability ^67^. Similar concepts have been proposed for PD ^161^. Our data cannot rule this scenario out and, notably, simulations over molecular and neurotransmission marker similarity networks did explain tau load better on average than simulations over anatomical networks.

Delving into the neurotransmission marker profile driving these analyses highlighted a prominent role of the serotonergic system, particularly 5-HT_1A_ and the ratio of 5-HT_1A_ and 5-HT_1B_ receptors. 5-HT_1A_ is critical for memory processes, is densely expressed in the medial temporal lobe, and is reduced in AD, which itself is associated with worse cognition ^162^. A PET study showed a strong relationship between 5-HT_1A_, clinical status, cognition and hippocampus atrophy, which was enhanced after correcting for brain atrophy, and was corroborated using autoradiography ^163^. Tau pathology has been described in the dorsal raphe nucleus (where 5-HT is synthesized), which is known to innervate the entorhinal cortex ^164^ and, interestingly, a recent study has described a reduction in tau pathology with plasma p-tau181 with selective serotonin reuptake inhibitors (SSRI) administration ^165^. 5-HT_1A_ receptors are also well understood to modulate excitatory-inhibitory (E:I) balance ^166,167^. Disrupted E:I balance has been highlighted as a key feature of AD, perhaps acting as a main driver of pathology accumulation and/or cognitive impairment ^168–173^. Our post hoc analysis found confluence between the spatial distribution of tau pathology and the ratio of excitatory and inhibitory receptors, supporting the notion that tau prone regions are characterized by a greater excitatory tone. However, this finding was restricted to ionotropic receptors, whereas the distribution ratio of metabotropic E:I receptors was anti-correlated with tau distribution. This distinction is important, since activation of metabotropic receptors results in slower and more long-lasting changes than does that of ionotropic receptors. Together, the results suggest that tau pathology gradually proceeds over a gradient of decreasing ionotropic excitatory potential, further highlighting the role of E:I imbalance in AD, and possibly highlighting an under-appreciated role of serotonergic modulation in these dynamics.

Subtyping analyses revealed that tau presence is highly consistent across individuals, whereas tau load within those regions shows significant inter-individual variability. Consistent with previous work ^28^, four tau load subtypes were identified. The Limbic and Lateral Temporal subtype patterns could both be explained by simulating tau spread through white matter fibers with regional *MAPT* expression modulating tau misfolding rate. The Posterior subtype tau pattern could also be recapitulated through anatomical spread of tau from an entorhinal epicenter, however our model search consistently nominated a role of acetylcholine (ACh) receptors and transporters in tau clearance for this subtype specifically. This is notable given that dysfunction of the ACh system has been thought of as an essential feature of AD, and given that acetylcholinesterase inhibitors are a common treatment for cognitive impairment due to AD ^174,175^. There is no current evidence that elevating ACh levels in the brain helps to reduce AD tau pathology, but there are also a dearth of studies investigating this in living humans. One study did find slower tau accumulation in patients taking acetylcholinesterase inhibitors ^165^, though this study warrants replication and spatial patterns have not been investigated. Future studies should probe the ACh systems of individuals with this AD subtype, and should investigate the effects of neuromodulating medication on tau patterning.

Finally, there is the MTL-sparing subtype, which has perhaps the most singular presentation and disease course. Besides its distinct cortical-predominant pattern, the MTL-sparing subtype consistently shows an earlier age of onset and a non-amnestic phenotype, shows less activated microglial/macrophage burden, lower likelihood of *APOE* e4 allele carriage, and more basal forebrain pathology ^28,122,176^. This has led to speculation that this subtype may have a unique etiology compared to other subtypes ^177^. In the present study, two of the top three models for the MTL-sparing subtype involved regional moderation based on *MAPT* and A*β*, again nominating traditional AD mechanisms. However, the pattern was most accurately reproduced when simulating tau over functional connectivity, with a precuneus epicenter (8 of top 10 models), pointing to a fairly unique model needed to reconstruct this pattern. The rapid accumulation of tau throughout the cerebral cortex in this subtype is almost reminiscent of A*β* accumulation, making an alternative spread mechanism for this subtype an interesting prospect. Spread of tau over functional networks is of course not compatible with an agent-based spread hypothesis, but is compatible with cascading network failure hypotheses ^48,50^. However, structural and functional connectivity approaches have known flaws, and each successfully captures aspects of known synaptic connectivity that the other cannot ^178^. Therefore, agent-based spread over white-matter connections cannot be ruled out in this case. With regard to a distinct epicenter in MTL-sparing cases, we did not find evidence for variation in the sequence of tau presence and, as noted, all MTL-sparing cases also feature some MTL pathology ^122,177^. In addition, well performing models did emerge using an entorhinal epicenter (with the top model also once again involving the 5HT_1A_ receptor), though they were not top performers. In all, these results continue the theme that a distinct etiology is possible in the MTL sparing subtype, a thesis that should continue to be further explored with future studies.

This study showcases the power and potential of mechanistic disease simulation models to both test and generate hypotheses from human data. This is important for a disease like AD that appears to be unique to humans. This type of *in silico* approach can be an important framework to build upon as new mechanisms are understood and, in turn, can help in directing the focus of future mechanistic studies. This study does come with several limitations that must be acknowledged. The SIR model fits the data under the assumption that different regional factors influence tau synthesis, misfolding, clearance, and spread. However, it is not a molecular kinetic model and assumes the same clearance rate for normal and misfolded tau, which may limit its ability to fully capture the complexity of tau pathology. Future studies should better integrate known tau kinetics as this information emerges. Additionally, most of the brain properties measured in this study are macroscale summaries of microscale phenomena. Some of these summaries are higher fidelity than others, and varying types of methodological error are likely present in all of our measures. In addition, using group averages for these factors may obscure individual variability that may be relevant for understanding tau progression. These factors make it difficult to conclude whether mechanistic interactions are actually taking place as the model suggests they are. Similarly, these interactions are based on spatial colocalization. While our simulations provide many possibilities for mechanistic interaction and test them against actual outcomes based on human tau-PET data, we cannot confirm causality without experimental validation. Some of the multimodal brain connectivity measurements and regional biological factors we used in the model do not include data for the hippocampus and amygdala—key regions in Alzheimer’s disease—and our exclusion of these structures potentially affected model performance and outcomes. The analyses are conducted on group average population tau-PET maps, and future work should work toward integrating person-specific models ^179,180^. Finally, while a great strength of our approach is that tau mechanisms are simulated dynamically, factors contributing to regional vulnerability in our model that are known to themselves be dynamic (e.g., A*β*, E:I imbalance) are nonetheless integrated in a static manner.

In conclusion, we describe possible mechanisms underlying two distinct processes of tau progression: Braak-like tau presence measured and tau load. While tau presence is primarily driven by connectome-based spreading and remains consistent across individuals, tau load is shaped by regional vulnerability, resulting in diverse patterns. Subtyping analyses further highlight the heterogeneity of tau load patterns and their unique drivers. Future studies should aim to validate these mechanisms in a patient specific and longitudinal context, while also identifying specific genetic factors that could inform therapeutic development. Such efforts will deepen our understanding of tau dynamics and support the advancement of tau-targeted, subtype-specific therapies.

## Methods

### Participants

Participants of this study represented a selection of individuals from the Swedish BioFINDER-2 Study (https://biofinder.se) (NCT03174938), which was designed to accelerate the discovery of biomarkers indicating progression of Alzheimer’s disease pathology ^181^. Participants for the present study were selected based on the following inclusion criteria: participants must (i) have a [^18^F]RO948-PET scan, (ii) be cognitively unimpaired, or classified as having mild cognitive impairment ^182^, or Alzheimer’s disease dementia ^183^, (iii) with biomarker evidence of *β*-amyloid (A*β*) positivity defined by CSF A*β*42/40 ratio obtained using the Elecsys immunoassays ^135^. The detailed inclusion and exclusion criteria was included in Supplementary Note <u>1</u>. Specifically, a pre-established cutoff of 0.08 for the CSF A*β*42/40 ratio, based on Gaussian mixture modeling ^181^, was used to define A*β* positivity. For participants without Elecsys measurements, clinical routine assays with pre-established cutoffs were used: Lumipulse G assay (cutoff of 0.072) ^184^ or Meso Scale Discovery assay (cutoff of 0.077). All participants fitting the inclusion criteria with [^18^F]RO948 scans acquired (BioFINDER-2) in March 2024 were included in this study. In total, 219 cognitively unimpaired, 212 mild cognitive impairment (MCI), and 215 suspected Alzheimer’s dementia (AD) individuals were included. Demographic information can be found in Supplementary Table S1. All subjects provided written informed consent to participate in the study according to the Declaration of Helsinki; ethical approval was given by the Ethics Committee of Lund University, Lund, Sweden, and all methods were carried out in accordance with the approved guidelines. Approval for PET imaging was obtained from the Swedish Medicines and Products Agency and the local Radiation Safety Committee at Skåne University Hospital, Sweden.

### PET acquisition and pre-processing

PET and Magnetic Resonance Imaging (MRI) acquisition procedures for BioFINDER-2 have been previously reported ^185^. Tau PET imaging was performed using [^18^F]RO948-PET, and all PET data were processed using a standardized volumetic pipeline described previously ^28,185^. Briefly, Pet data were reconstructed into four 5-min frames acquired between 80 and 100 min post-injection. Frames were re-aligned using AFNI’s *3dvolreg* (https://afni.nimh.nih.gov) and averaged within participant, and rigidly coregistered to each subject’s native space T1-weighted MRI image. The coregistered image was intensity-normalized using an inferior cerebellar gray reference region, creating standard uptake value ratios (SUVR) iamges. T1-weighted MRI images were preprocessed using FreeSurfer v6.0 (https://surfer.nmr.mgh.harvard.edu) to obtain native-space cortical and subcortical parcellations based on the Desikan-Killiany (DK) atlas ^80^. Regional mean SUVR values were extracted within these anatomically defined regions in native space. To obtain the group-level regional map of A*β*, [^18^F]-flutemetamol-PET images were preprocessed for A*β* positive subjects with subjective cognitive decline, MCI and AD. SUVRs were calculated using the cerebellar cortex as the reference region and the group-level regional map was derived by averaging the A*β* SUVRs across these participants ^186^. Additional details on PET acquisition, image reconstruction, preprocessing parameters, and diagnostic criteria are provided in the Supplementary Note <u>2</u>.

### Regional tau-PET data preprocessing

#### Tau Load (SUVR)

Preprocessing of [^18^F]RO948-PET data yielded mean regional tau-PET SUVR values, extracted from each participant’s native space PET image using the FreeSurfer-derived DK atlas ^80^. Analysis focused exclusively on cortical regions, hippocampus and amygdala, resulting in 66 regions in total ^72^. To accurately represent tau load, regional SUVR values were MinMax normalized using a reference group of A*β*-negative, cognitively normal elderly participants, with adjustments for choroid plexus binding ^28,72^. This normalization helps keep values in the expected range of the SIR model, and regional normalization to a reference group helps to mitigate regional perfusion and off-target binding patterns, the latter of which has been a well-documented issue in [^18^F]RO948-PET studies ^187^ and can lead to erroneous signals in regions not accumulating tau pathology.

#### Tau presence (TPP)

Tau-positive probability (TPP) was calculated to represent tau presence. For each brain region, SUVR values were processed using a two-component Gaussian mixture model (GMM), based on the assumption that tauopathy-related and tauopathy unrelated signals across the population form distinct Gaussian distributions ^72^. The posterior probability from this GMM was used to determine the likelihood that a participant’s SUVR fell within the tau-positive distribution, providing a probabilistic measure of tau presence in each region.

To investigate differences between tau load (SUVR) and tau presence (TPP), we examined their relationships with Braak stages, tau-PET SUVR, and CSF biomarkers. Brain regions were first grouped into six Braak stages ^19^, and the two tau measures (SUVR and TPP) were compared across these stages. For each participant, regional tau measurements were categorized into three Braak stage groups: early (I), intermediate (III/IV), and late (V/VI). Within each group, tau-PET SUVR was compared to min-max tau presence and tau load values. Lastly, the relationships of CSF A*β*42/40 ratio and CSF p-tau217 with min-max normalized tau load and presence values were analyzed across the three Braak stage groups.

### The Susceptible-Infectious-Recovered Model

To investigate mechanisms of tau propagation, we employed the Susceptible-Infected Removed (SIR) agent-based model to simulate tau load and distribution ^77^. This connectome-based disease spreading model assumes that tau pathology originates from a predefined disease epicenter, propagating through the brain along the connections, while being modulated by the intrinsic properties of each brain region. This approach enables exploration of the interplay between connectome-based spreading and regional vulnerability in shaping tau propagation.

Briefly, the model simulates four coupled processes governing tau dynamics over time: spread synthesis, clearance, and misfolding (Fig. 2a). At each simulation step, tau propagates along the structural connectome, new normal tau is synthesized, existing tau is cleared, and a fraction of normal tau converts into misfolded tau. Model outputs were compared against observed regional tau presence (TPP) and tau load (SUVR).

Each process is governed by a combination of global hyperparameters that are constant across regions (Fig. 2a grey boxes) and region-specific parameters that encode intrinsic regional vulnerability (Fig. 2a red boxes). Global parameters regulate overall tau spreading velocity, misfolding probability, and retention of tau within neuronal compartments, whereas regional parameters modulate the local (regional) rates of spread, synthesis, clearance, and misfolding. All processes are sequential and depend on the protein levels from the previous simulation step, scaled by the simulation time increment (*Δt*). Further details regarding the model’s equations and formulation are provided in Supplementary <u>E1</u>-<u>E6</u> and ^77,78^.

The original SIR formulation includes regional modulation of tau synthesis and clearance^77,78^. To better capture the biological complexity of tau propagation, we extended the original model to additionally incorporate region-specific modulation of tau spread rate (Supplementary <u>E1</u>) and misfolding rate (Supplementary <u>E4</u>).

Model hyperparameters were tuned to optimize correspondence between simulated and observed tau patterns. Tested parameters included tau retention probability, global misfolding rate, and spreading velocity (see Supplementary <u>N3</u> for hyperparameter ranges). Full mathematical formulation of the model, including governing equations and variable definitions, is provided in Supplementary <u>E1</u>-<u>E6</u>.

### Alternative connetome-based spreading models

To assess whether the dissociation between tau presence and tau load was specific to the SIR framework, we repeated key analyses using two independent, previously established approaches. First, we applied the Epidemic Spreading Model (ESM), a diffusion-based model that simulates signal propagation along the structural connectome from a predefined epicenter ^72,83^. Using the entorhinal cortex as the epicenter, consistent with neuropathological evidence and Extended Data Fig. 2, ESM predictions were compared against observed regional tau presence (TPP) and tau load (SUVR). Second, we applied the connectivity-based statistical approach introduced by *Franzmeier et al., 2020*, relating regional tau spreading sequences to connectivity-derived distances from empirically defined epicenters, as described previously ^52,84,85^. For both approaches, we used the same 66-region weighted structural connectivity (SC) matrix as in Fig.2, derived from a young, healthy cohort using deterministic tractography from DWI data ^72,188^ (see Structural Connectivity). Full methodological details are provided in the Supplementary Note <u>4</u>, <u>5</u>.

### Conventional AD-related elements

To investigate the influence of conventional AD-related elements on tau propagation, we first tested the trans-neuronal spread hypothesis. Based on our previous work ^72^, we used a low-resolution, 66-region weighted structural connectivity (SC) network derived from a young, healthy cohort using deterministic tractography from DWI data ^72,188^ (see Structural Connectivity). Tau propagation was simulated from a bilateral entorhinal cortex seed region (epicenter) throughout the anatomical connections (i.e. SC) across the 66 regions, with all four mechanistic rates (synthesis, clearance, spread, misfolding) set uniformly across regions. The simulation was done across 30,000 time steps. Pearson correlations were computed at each time point between simulated and empirical tau patterns to evaluate model fit, and the simulated pattern showing the highest correlation with empirical data at a given time point was selected as the final tau pattern for analysis. Additionally, the model performance was tested using each brain region as a potential epicenter to determine the optimal initiation site for tau propagation.

We next tested whether regional vulnerability interacts with brain connectivity by incorporating three well-established AD-related factors (A*β* deposition, *APOE*, and *MAPT* gene expression) as input parameters to modulate the mechanistic rates of tau propagation in the trans-neuronal spread model. Each regional factor was parameterized to influence one of the four mechanistic rates: synthesis, clearance, spread, or misfolding. For each regional factor, we selected the model with the best performance (highest Pearson correlation coefficient R-value) among the four possible propagation mechanisms, which was then used for subsequent analyses. The selected mechanism was interpreted as the most likely pathway by which the regional factor influences tau propagation. All models were run separately for tau load (SUVR) and tau presence (TPP).

### Testing alternative hypotheses of tau propagation

To explore other possible factors influencing tau propagation, we assessed the influence of various brain connectomes and regional biological factors by substituting the anatomical connectivity and conventional AD-related factors with novel brain connectomes and regional factors (see Novel brain connectivity measurements and Novel biological factors sections below). Different connectomes allowed us to investigate alternative hypotheses of tau spread, while the inclusion of diverse regional factors provided insights into intrinsic properties that might shape tau presence and load patterns. By analyzing potential interactions between these elements, we aimed to identify whether alternative combinations of connectivity and regional factors could better explain tau propagation compared to known or conventional factors.

A total of 1,576 models were evaluated, including eight connectome-only models, and combinations of eight connectomes with 49 regional factors, each modulating four tau propagation mechanisms: spread, synthesis, misfolding, and clearance. Only models meeting the following criteria were selected for further analysis: (i) connectome only models, with all four mechanistic rates uniformly set across regions; (ii) the top performing connectome-regional factor models, determined by identifying the highest performing model among the four possible propagation mechanisms; and (iii) models that performed better than the 95% confidence interval of the null distribution (see Statistical analysis section below). The analyses were restricted to 62 cortical regions, excluding the hippocampus and amygdala, as these regions were unavailable for certain connectomes. Separate analyses were conducted for tau load (SUVR), tau presence (TPP), and the four identified SUVR subtypes (see SuStaIn section below).

### Brain connectome measurements

#### Structural connectivity

To test the trans-neuronal spread hypothesis, we constructed SC from 60 young healthy subjects (18-60 yrs, 33.08*±*12.54) from the CMU-60 DSI Template ^188^ to represent anatomical brain connections. The data processing pipeline has been previously described in detail ^72^. Briefly, orientation distribution functions were computed and subsequently used to generate deterministic tractography between brain regions defined by the Desikan-Killiany atlas. The resulting connectomes were averaged across all participants, yielding a template SC network comprising 66 regions of interest, representing a healthy population. We also tested an alternative young SC network derived from 62 cortical regions, excluding the hippocampus and amygdala, based on data from 326 young participants (ages 22–35) in the Human Connectome Project (HCP) ^87,88^.

#### Multimodal brain networks

We utilized eight distinct cortical connectomes ^87^ to test various hypotheses of tau propagation (Fig. 3a). These connectomes were derived from diverse modalities, each reflecting different biological and functional properties: SC from diffusion tractography to model direct anatomical pathways; functional connectivity (FC) from functional MRI (fMRI)^88,189^, electrophysiological signal covariance (EP) from Magnetoencephalography (MEG) ^190,191^, and temporal profile similarity (TS) across fMRI-derived timeseries to explore the hypothesis of tau spreading across actively communicating neurons ^192–194^; gene co-expression covariance (GC) across 20,000 genes to investigate cascading vulnerability due to shared gene expression ^86,195^; neurotransmission marker similarity (NS) from PET to examine a mixed hypothesis incorporating trans-synaptic propagation and neurotransmission-driven vulnerability ^89,90^; glucose metabolism covariance (MC) from Fluorodeoxyglucose-PET to test whether glucose metabolism contributions to tau spread ^196^; laminar similarity (LS) from postmortem brain to assess evolutionary and developmental influences on tau vulnerability ^197–199^.

In further detail, SC reflects the probability and the strength of white matter connections between two brain regions, while FC captures the synchronization of resting-state BOLD activity. Both SC and FC data were obtained from 326 healthy participants (ages 22–35) from the HCP ^88^. TS, calculated from statistical fMRI time series features, measures the similarity of temporal dynamics between cortical regions across over 7,000 local features. RC, based on PET imaging of 18 neurotransmitter receptors and transporters, indexes neurotransmission marker density similarity between regions, covering dopamine, serotonin, acetylcholine, norepinephrine, histamin, cannabinoid, opioid, glutamate and GABA systems ^90^. GC reflects transcriptional similarity between cortical regions, derived from six postmortem brains (ages 24–57 years) from the Allen Human Brain Atlas ^86^ (AHBA; http://human.brain-map.org/). LS, estimated from BigBrain histological data (65-year-old male) ^198^, captures similarity in cellular distributions across cortical layers. MC, based on [^18^F]-fluorodeoxyglucose PET from 26 healthy participants (77% female, ages 18–23) ^196^, represents similarity in glucose metabolism between brain regions. EP, obtained from MEG recordings of 33 participants (ages 22–35) from HCP ^88^, assesses connectivity via magnetic fields produced by neural currents. Further details on data acquisition and preprocessing have been previously detailed ^87^ and visualized in Fig. 3a.

All connectomes were normalized using the arctanh transformation ^87^ and restricted to positive values. All connectomes were constructed for the 62 cortical regions, derived by excluding the hippocampus and amygdala from the original set of 66 regions, using the DK atlas. For FC, the top 40% of positive connections were retained for each participant to emphasize the strongest associations, and the connectomes were then averaged across participants. Thresholds of 10%, 20%, 30%, and 40% were tested, with only the 40% threshold resulting in average connectivity without isolated regions.

### Regional biological factors

#### Conventional AD-related factors - Aβ, MAPT, APOE

To assess whether regional vulnerability interacts with brain connectivity to influence tau propagation, we first evaluated three well-established AD-related factors: regional A*β* deposition, *MAPT* and *APOE* gene expression. Regional A*β* deposition was approximated using group-averaged A*β* SUVR values of 66 regions of interest from the BioFINDER-2 dataset. *MAPT* and *APOE* gene expression data were extracted from the AHBA ^86^, which provides regional microarray expression data from six post-mortem brains (one female, ages 24–57 years, 42.5 ± 13.38 years). AHBA data were preprocessed and mapped to the brain regions of interest using the abagen toolbox ^195^, and the final regional expression profiles were averaged across donors. In short, gene expression scores were generated for each of the 33 cortical parcels for both hemispheres, with data mirrored across hemispheres to increase spatial coverage, resulting in gene expression profiles for 66 regions. These regional factors were converted to probabilities by applying the cumulative distribution function to their z-score normalized values, and incorporated into the SIR model to regulate tau synthesis, clearance, spread, and misfolding rates. Greater standardized regional factor probability corresponded to higher rates for these processes.

#### Novel biological factors

To explore other intrinsic properties that might shape tau patterns, we tested 49 novel regional biological factors (Extended Data Fig. 4): 19 neurotransmitter receptors and transporters ^90^, six measures of cortical expansion ^89^, four metabolic features ^89^, seven cell-type profiles ^91^, three structural properties ^89,91^, and ten AD-related factors, including seven gene clusters (Extended Data Fig. 6) identified from large GWAS studies ^92–94^ (see below) along with *MAPT*, *APOE*, and A*β*.

The neuromaps toolbox ^89^ (https://github.com/netneurolab/neuromaps) was used to obtain regional maps for all factors except the AD-related factors. Further details about the acquisition and processing of these maps are available in ^89,91^. To extract AD-related gene clusters, we identified 90 genes from three recent large GWAS studies, 79 of which are available in the AHBA dataset (Supplementary Table S7). Regional expression maps of the available 79 genes were processed using the abagen toolbox. Given that many genes show fairly consistent expression patterns, we opted to summarize genes with similar expression patterns using hierarchical clustering. Seven clusters provided the best fit (tested from 2 to 10 clusters) and were used in subsequent analyses (Extended Data Fig. 8). The similarity among 49 regional factors is shown in Extended Data Fig. 8.

### Neurotransmission marker properties

We evaluated the similarity of each of the seven brain connectomes (Fig. 3a) to the SC network using three metrics: structural similarity index measure (SSIM), cosine similarity, and Pearson correlation (R). SSIM emphasizes structural consistency by accounting for spatial organization, cosine similarity measures the angular relationship between vectors, and R quantifies the linear correlation.

To understand why the neurotransmission marker similarity network best recapitulates tau presence/load, we assessed correlations between neurotransmission markers distributions and tau load- /distribution patterns using Pearson correlation coefficient (R) and examined neurotransmission marker density variations across six Braak stages. Each of the 19 individual neurotransmission markers (15 receptors and 4 transporters) was analyzed, alongside “a priori” and “a posteriori” neurotransmission marker ratios grouped based on existing literature and our findings, respectively (Supplementary Table S8).

The “a priori” neurotransmission marker combinations reflected established neurotransmission functions. Five excitatory to inhibitory ratios were calculated: 1) the overall ratio ***E:I***, capturing all neurotransmitters, calculated as *(5-HT_2A_ + 5-HT_4_ + 5-HT_6_ + D_1_ + M_1_ + α_4_β_2_ + NMDA + mGluR_5_) / (5-HT_1A_ + 5-HT_1B_ + D_2_ + GABA_A/BZ_ + H_3_)*; 2) the E:I ratio specific to metabotropic neuroreceptors ***E:I_metabotropic_***, was defined as *(5-HT_2A_ + 5-HT_4_ + 5-HT_6_ + D_1_ + M_1_ + mGluR_5_) / (5-HT_1A_ + 5-HT_1B_ + D_2_ + H_3_)*; 3) the E:I ratio specific to ionotropic neuroreceptors ***E:I_iontropic_***, was defined as *(α_4_β_2_ + NMDA) / GABA_A/BZ_;* 4) the glutamate-to-GABA ratio ***E:I_Glu/GABA_***, calculated as *(NMDA + mGluR_5_) / GABA_A/BZ_*; 5) the iontropic glutamate-to-GABA ratio ***E:I_Glu/GABA(ion)_***, was defined as *NMDA / GABA_A/BZ_*. Additionally, the ratio of receptor-to-reuptake specific to acetylcholine ***R/Re_ACh_***was calculated as *`(M1 +` α_4_β_2_) / VAChT*, and the ratio of traditional neurotransmitters to neuromodulator ***NT/NM*** was calculated as *(GABA_A/BZ_ + mGluR_5_ + NMDA) / (5-HT_1A_ + 5-HT_1B_ + 5-HT_2A_ + 5-HT_4_ + 5-HT_6_ + A4B2 + D_1_ + D_2_ + M_1_ + H_3_)*.

The “a posteriori” neurotransmission marker combinations were derived from our observations of top-contributing neurotransmission markers in this study. These ratios have no known biological meaning or justification, but were rather “data-driven” combinations stemming from data in this study. They should therefore be interpreted with caution. These included the serotonin receptor ratio ***5-HT_1A_/5-HT_1B_***, representing serotonin receptor functions, and the dopamine-to-norepinephrine transporter ratio ***DAT/NET***, highlighting the balance between dopaminergic and noradrenergic activity.

### Analysis of tau subtypes

To determine whether tau load and distribution result in distinct subtypes, we performed de novo the Subtype and Stage Inference algorithm (SuStaIn) ^200^ on 1,475 [^18^F]RO948-PET images (Including A*β*-cognitive unimpaired participants) to identify distinct spatiotemporal trajectories of tau propagation. As in previous work ^28^, A*β*-negative cognitively unimpaired participants were specifically included in this analysis. SuStaIn combines disease progression modeling with clustering to infer multiple tau propagation patterns from cross-sectional data, providing both probabilistic and individualized classification of tau trajectories. The model uses spatial features (e.g., brain regions) and pseudo-temporal severity markers (e.g., Z-scores) to describe linear biomarker changes across disease stages, resulting in distinct subtype trajectories representing different sequences of tau pathology.

Following the established methodology ^28^, we utilized ten spatial features, including bilateral parietal, frontal, occipital, temporal, and medial temporal lobe regions, along with severity cutoffs derived from standardized tau levels. The number of subtypes (i.e., distinct spatiotemporal progressions) was determined via cross-validation. For each of k=1–4 subtypes, ten fold cross-validation was performed by fitting SuStaIn to 90% of the data and evaluating sample likelihood on the remaining 10%. Model fit was assessed using cross-validation-based out-of-sample log-likelihood ^200^. Separate analyses were conducted for tau load and tau presence. For tau load (SUVR), the log-likelihood increased consistently up to k = 4, suggesting the presence of at least four distinct subtypes. For tau presence (TPP), however, the log-likelihood improved only up to k = 2, with no additional benefit observed beyond this point. As the identification of a second subtype for tau presence seemed to be influenced by model uncertainty and hemispheric laterality, we repeated the subtyping analysis using five spatial features by combining left and right regions. This revised approach indicated that the optimal number of subtypes for tau presence was k = 1 (Fig. S2b).

Next, we sought to examine the connectome and regional features contributing to distinct tau subtypes. To ensure consistency, we aimed to stay as close to the original subtypes as possible, using the previously published ones for this analysis ^28^. We extracted the average tau load (SUVR) across participants for each subtype, based on the original participant list from the BioFINDER subset used in the prior study. This approach was used to leverage the robustness of previously validated subtypes, which were confirmed across multiple cohorts and different tau-PET tracers. The four subtypes extracted from the prior work were consistent with those identified by the SuStaIn model in the current analysis (Fig. S2c). Subsequently, we employed SIR models to investigate individual variability in tau propagation by testing alternative hypotheses of tau propagation on each of the four subtype patterns separately. Additionally, we evaluated the influence of different epicenters. Specifically, our previous work suggested early-stage epicenters of each subtype fell in either the medial temporal lobe or precuneus ^28^, so we additionally tested the effect of entorhinal cortex or precuneus on tau spread within each subtype.

### Statistical analysis

The SIR was applied using various brain connectomes and regional factor combinations (see above). Each model was assessed for fit across the entire sample of A*β*-positive individuals, quantified using the Pearson correlation coefficient (R) between simulated and observed regional tau patterns. Specifically, Pearson correlations were calculated at each simulated time point to capture changes in model performance throughout the simulation. Model fit was defined as the highest R value achieved across the entire simulation period, representing the best alignment between simulated tau and empirical data under the given hyperparameters and input data. Each model was independently evaluated with different sets of hyperparameters, and the configuration yielding the highest model fit was selected as the optimal configuration for the corresponding input connectome and regional biological factors.

To determine whether model performance exceeded chance levels, select models were compared against a null distribution generated from 1,000 surrogate brain connectome matrices with preserved degree and strength distributions, using the Brain Connectivity Toolbox (https://sites.google.com/site/bctnet/). Each null connectome matrix was input into the SIR model to simulate tau progression, with hyperparameters tuned to obtain the best fit. The Pearson correlation coefficient (R) values for all 1,000 null matrices were used to create a null distribution, from which the mean and 95% confidence intervals were calculated to determine the likelihood of the observed performance occurring by chance. This analysis was repeated separately for the eight input brain connectomes and for simulations involving tau load (SUVR), tau presence (TPP), and each of the four SUVR subtypes.

In addition to the R-value, we evaluated the performance of the selected models using mean squared error (MSE) and explained variance (EV). MSE quantifies the average squared difference between observed and simulated tau pattern, whereas EV measures the proportion of variance in the observed tau explained by the model.

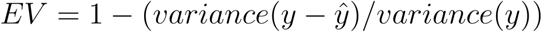

## Data availability

The structural and functional connectomes used in this study are openly available from ^87^ and can be accessed via GitHub (https://github.com/netneurolab/hansen_manynetworks/tree/v1.0.0). The neuromaps software toolbox, utilized for brain mapping analyses, is available at https://github.com/netneurolab/neuromaps, with documentation provided on GitHub Pages (https://netneurolab.github.io/neuromaps/). Further details can be found in the associated publication ^89^. Cortical layer thickness data are publicly available through BigBrainWarp (https://bigbrainwarp.readthedocs.io/en/latest/). Gene expression data used in this study were obtained from the Allen Human Brain Atlas and are openly accessible at https://human.brain-map.org.

The BioFINDER data are not publicly available, but access requests of anonymized data can be made to the study’s steering group. Access to the data will be granted in compliance with European Union legislation on the General Data Protection Regulation (GDPR) and decisions by the Ethical Review Board of Sweden and Region Skåne. Data transfer will be regulated under a material transfer agreement.

## Code availability

Python3.9 was used to run all models and analyses. Scripts for the revised Susceptible-Infectious-Recovery model can be found at https://github.com/DeMONLab-BioFINDER/Xiao_Tau_simulation_SIR. Implementations of the SuStaIn algorithm are available on the UCL-POND GitHub page: https://github.com/ucl-pond.

## Acknowledgements

This work was supported by the SciLifeLab Wallenberg Data Driven Life Science Program (grant: KAW 2020.0239), the National Institute of Aging (U01 AG079847-02), Swedish Alzheimer Foundation (AF-994626), and the Swedish Research Council (2024-03642). The BioFINDER-2 study was funded by the National Institute of Aging (R01AG083740), the European Research Council (ADG-101096455), the Swedish Research Council (2022-00775), Alzheimer’s Association (ZEN24-1069572, SG-23-1061717), GHR Foundation, ERA PerMed (ERAPERMED2021-184), Knut and Alice Wallenberg foundation (2022-0231), Strategic Research Area MultiPark (Multidisciplinary Research in Parkinson’s disease) at Lund University, Swedish Brain Foundation (FO2021-0293, FO2023-0163), Parkinson foundation of Sweden (1412/22), Swedish Alzheimer Foundation (AF-980907, AF-994229), Cure Alzheimer’s fund, Rönström Family Foundation, Konung Gustaf V:s och Drottning Victorias Frimurarestiftelse, Skåne University Hospital Foundation (2020-O000028), Regionalt Forskningsstöd (2022-1259) and Swedish federal government under the ALF agreement (2022-Projekt0080, 2022-Projekt0107). The precursor of 18F-flutemetamol was sponsored by GE Healthcare. The precursor of 18F-RO948 was provided by Roche. The funding sources had no role in the design and conduct of the study; in the collection, analysis, interpretation of the data; or in the preparation, review, or approval of the manuscript.

## DISCLOSURES

OH is an employee of Lund University and Eli Lilly.

NMC has received consultancy/speaker fees from Biogen, Eli Lilly, Owkin, and Merck.

GS has received consultancy/speaker fees from Springer, Adium, and GE. JWV has received consultancy/speaker fees from Manifest Technologies

## Extended data

**Extended Data Fig. 1.**
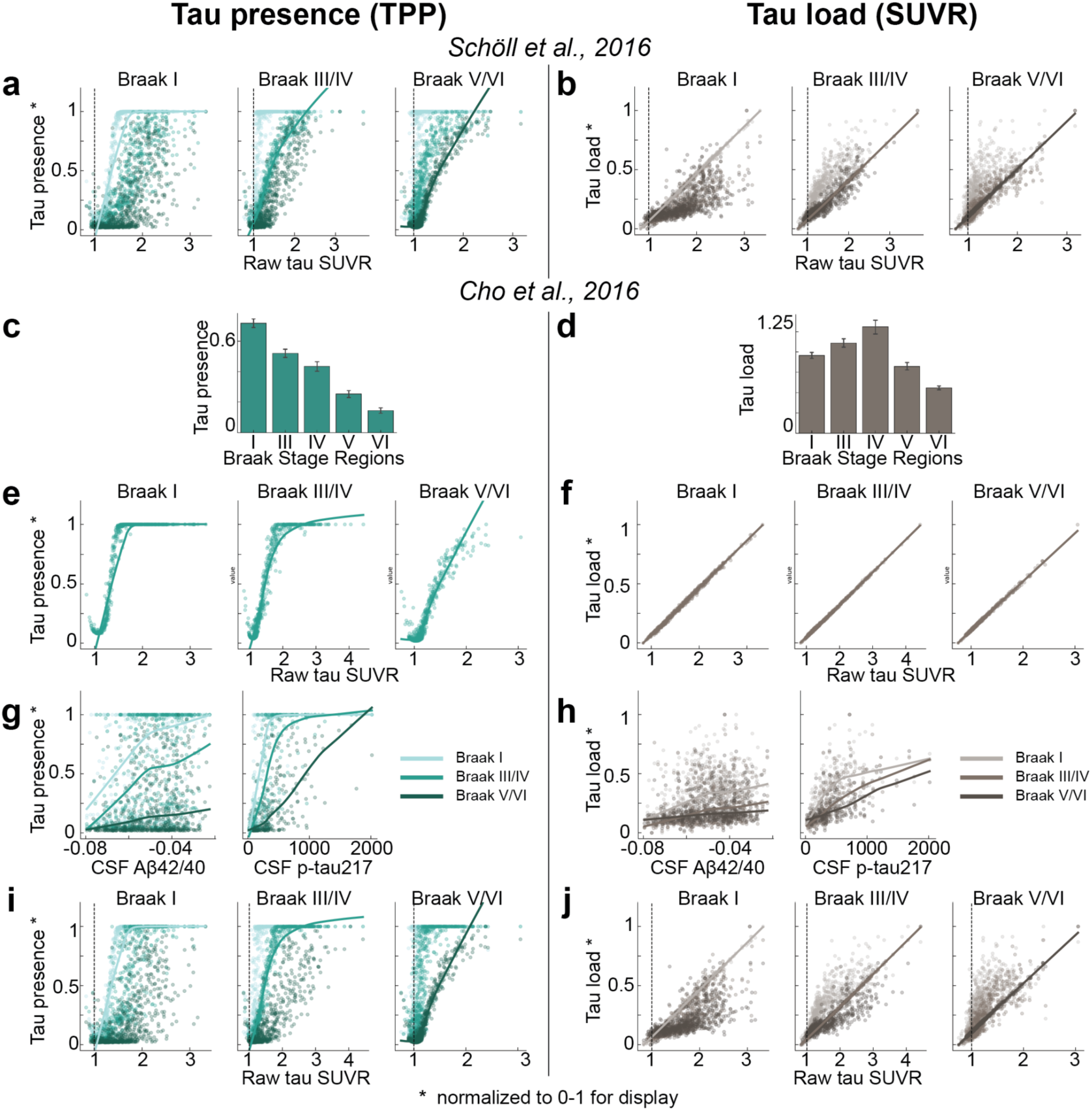
Relationship of tau presence and tau load with raw SUVR values. Related to Fig. 1. **a, b)** Relationship between tau presence a) and tau load b) and raw SUVR values across regions grouped by Braak stages, as defined by ^19^. In each panel, the fitted line is estimated using only regions belonging to the indicated Braak stage, while regions from other Braak stages are shown in different colors for comparison. **c-j)** Relationship between tau presence/load with Braak stage regions derived from ^20^. **c, d)** mean ± SEM values of tau presence c) and load d) for Braak stage regions. **d, f)** Relationship of tau presence e) and tau load f) with raw SUVR values in grouped regions of ^20^ Braak stages. **g, h)** Relationship of tau presence g) and load h) with CSF A*β*42/40 and CSF p-tau217 levels. **i, j)** Relationship between tau presence (I) and tau load (J) and raw SUVR values across regions grouped by Braak stages, as defined by ^20^. Regions corresponding to other Braak stages are shown in different colors for comparison. SUVR: standardized uptake value ratio; TPP: tau-positive probability; CSF: cerebral spinal fluid; Aβ: beta-amyloid; p-tau: phosporalated tau.

**Extended Data Fig. 2.**
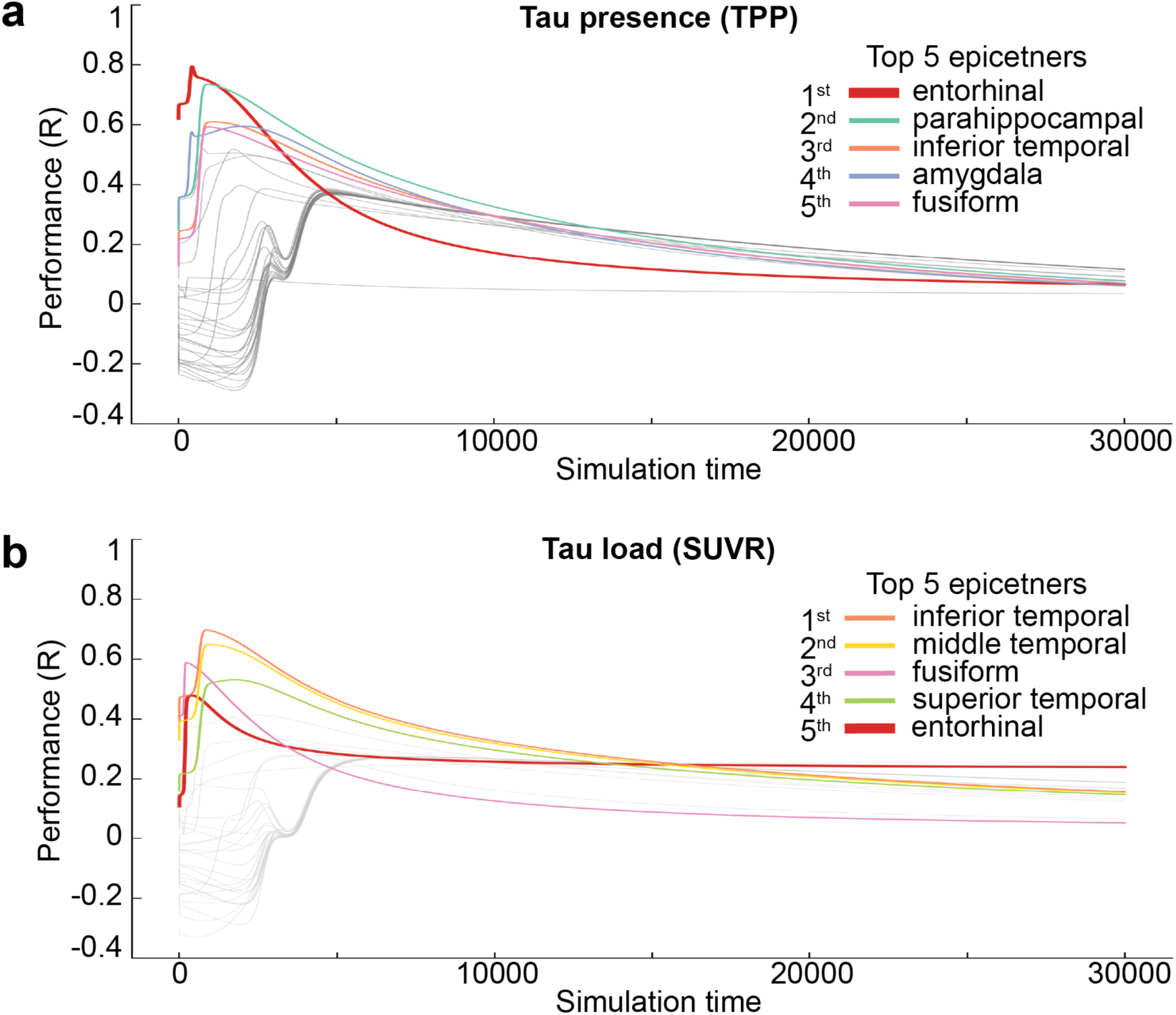
Model performance across time by varying epicenters. Related to Fig. 2. The spreading process was initiated in every brain region and the correlation between the simulated and empirical patterns of TPP **a)** and SUVR **b)** was computed across 30,000 simulation times. The five best seeding regions were highlighted. TPP: tau-positive probability; SUVR: standardized uptake ratio; R: Pearson correlation coefficient.

**Extended Data Fig. 3.**
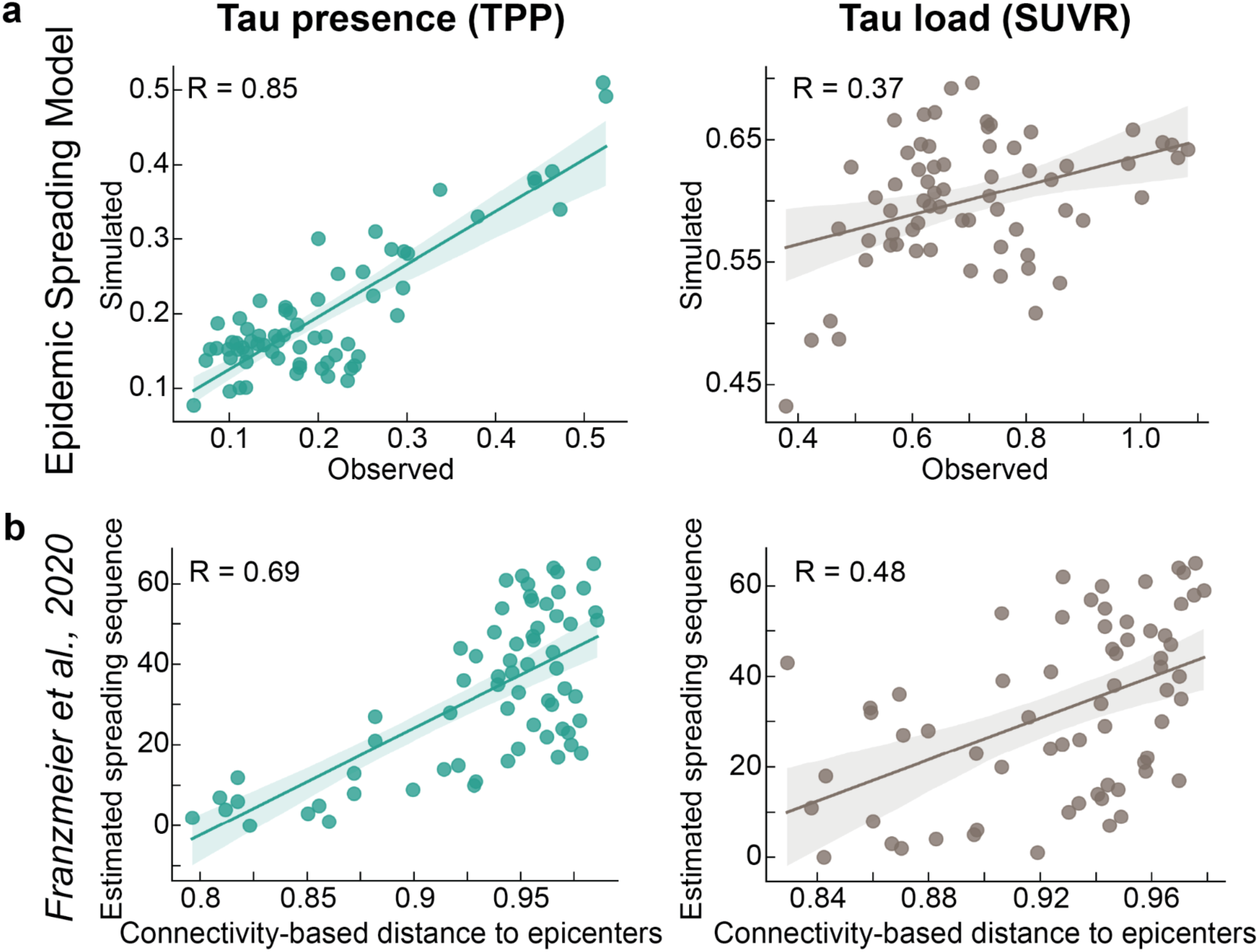
Using two different models, anatomical connections captures patterns of tau presence more than load. Related to Fig. 2. **a)** ESM model performance for tau presence (left) and tau load (right). ESM model simulates tau diffusion through the structural connectome and use the entorhinal cortex as epicenter, same as SIR model in Fig.2a,2c. Scatter plots compare ESM model predictions to observed tau presence/load. **b)** Relationship between network connectivity and regional tau levels using inferential statistics. Scatter plots illustrate the association between connectivity-based distance to the epicenters and the tau positivity sequences for tau presence (left) and load (right). ESM: Epidemic spreading model; SUVR: standardized uptake value ratio; TPP: tau positive-probability.

**Extended Data Fig. 4.**
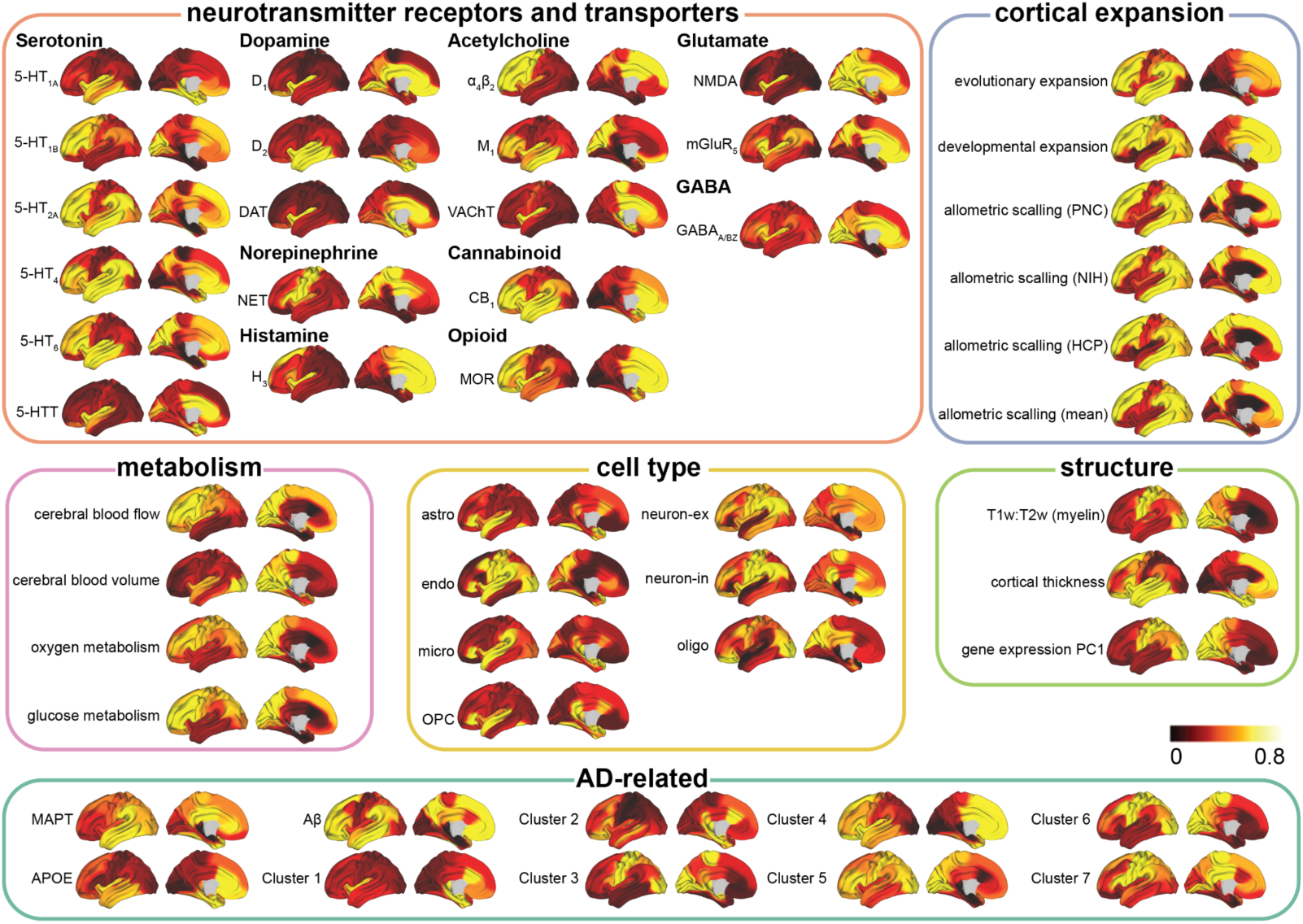
Cortical surface projections of input regional biological factors. Related to Fig. 3. neuromaps toolbox, the Allen Human Brain Atlas (AHBA) and the abagen toolbox ^195^ were used to compile a set of 49 micro-architectural brain maps, including measures of receptors and transporters, cortical expansion, metabolism, cell type-specific gene expression, microstructure, and AD-related factors (conventional: *MAPT*, *APOE*, A*β*; and seven derived AD-related gene clusters) (see *Methods* for more details). astro = astrocytes; endo = endothelial cells; micro = microglia; neuron-ex = excitatory neurons; neuron-in = inhibitory neurons; oligo = oligodendrocytes; and opc = oligodendrocyte precursors.

**Extended Data Fig. 5.**
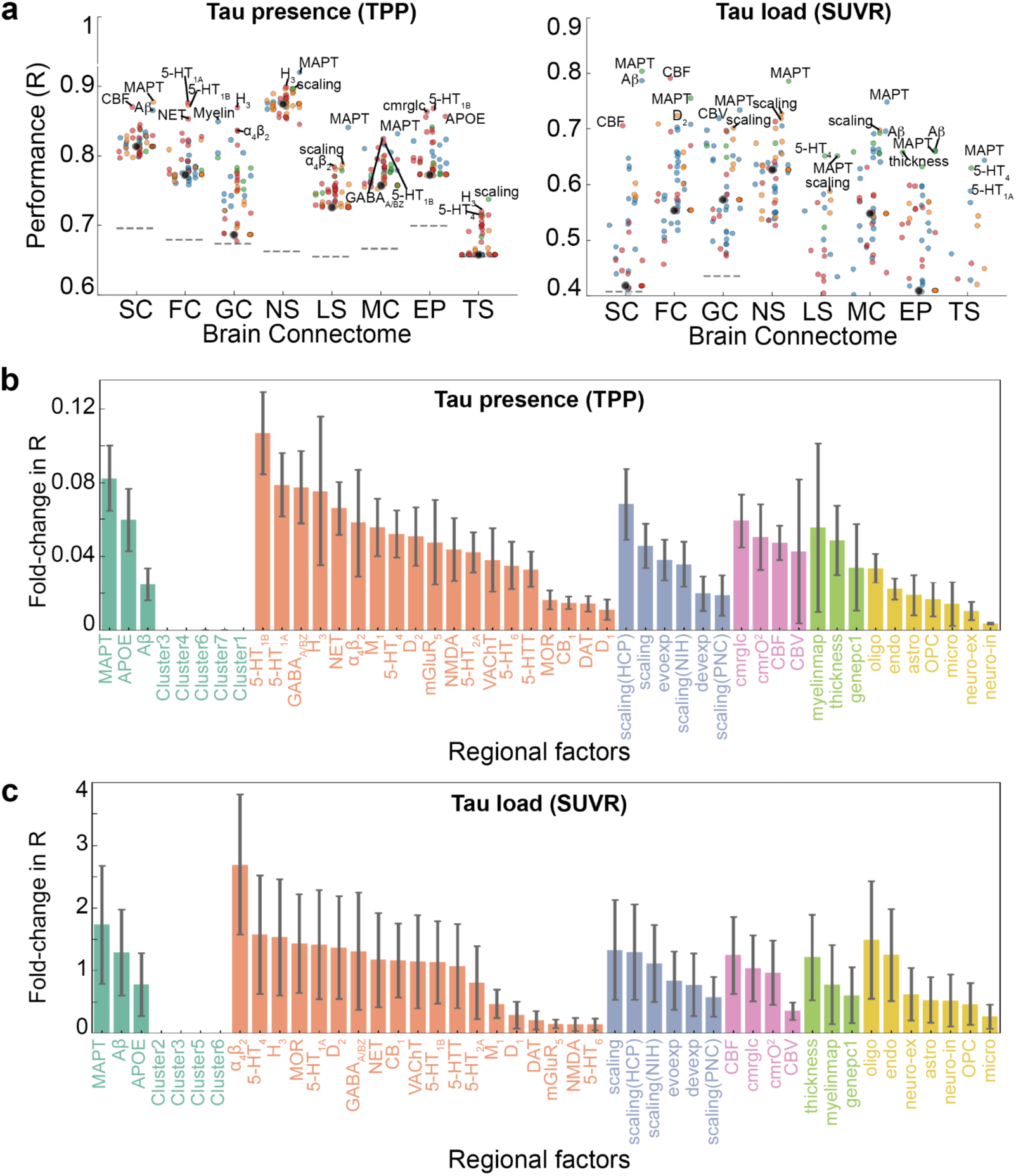
Model performance and contribution of regional biological factors over connectome only models for tau presence and load. Related to Fig. 3. **a)** Performance of individual models fitted to tau presence (TPP, left) and load (SUVR, right), respectively. This panel provides a zoomed-in view of *Main Text* Fig. 3e and 3f, with the y-axis range adjusted to 0.6-1 for tau presence and 0.4-0.9 for tau load. Each dot represents one model. The dot color indicates what parameter the regional information was influencing. Grayed-out dots did not exceed null model performance (indicated by dashed lines). The regional factor added to the model is labeled for the top 3 models of each connectome. **b, c)** Contribution of regional biological factors over connectome only models for tau presence b) and load c). Performance improvement over connectome only models was calculated and summarized by the biological factors shown on the x-axis, with results color-coded by the six types of regional biological factors. Regional factors for which models outperformed chance and the connectome-only model are shown (outperform chance: 95% CI of 1000 models scrambling connectomes but preserving weight and degree distribution). TPP: tau-positive probability; SUVR: standardized uptake ratio; SC: structural connectivity; FC: functional connectivity; GC: gene coexpression; NS: neurotransmission marker similarity; LS: laminar similarity; MC: metabolic covariance; EP: electrophysiological; TS: temporal similarity, R: Pearson Correlation coefficient R; astro = astrocytes; endo = endothelial cells; micro = microglia; neuron-ex = excitatory neurons; neuron-in = inhibitory neurons; oligo = oligodendrocytes; and opc = oligodendrocyte precursor cells.

**Extended Data Fig. 6.**
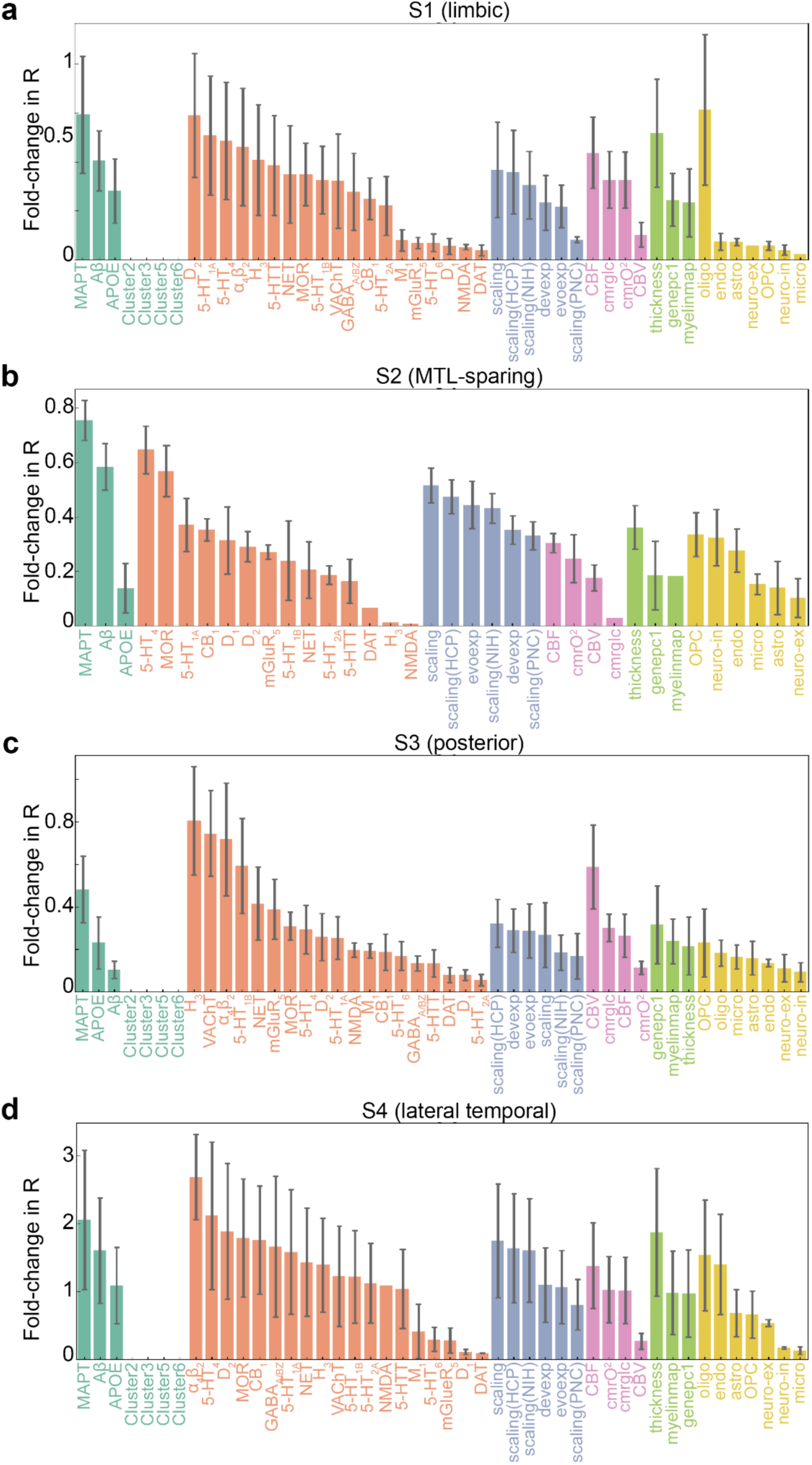
Contribution of regional biological factors over connectome only models for four tau load subtypes. Related to Fig. 5. Performance improvement over connectome only models was calculated and summarized by the biological factors shown on the x-axis, with results color-coded by the six types of regional biological factors. Regional factors for which models outperformed chance and the connectome-only model are shown (outperform chance: 95% CI of 1,000 models scrambling connectomes but preserving weight and degree distribution). MTL: medial temporal lobe; astro = astrocytes; endo = endothelial cells; micro = microglia; neuron-ex = excitatory neurons; neuron-in = inhibitory neurons; oligo = oligodendrocytes; and opc = oligodendrocyte precursor cells.

**Extended Data Fig. 7.**
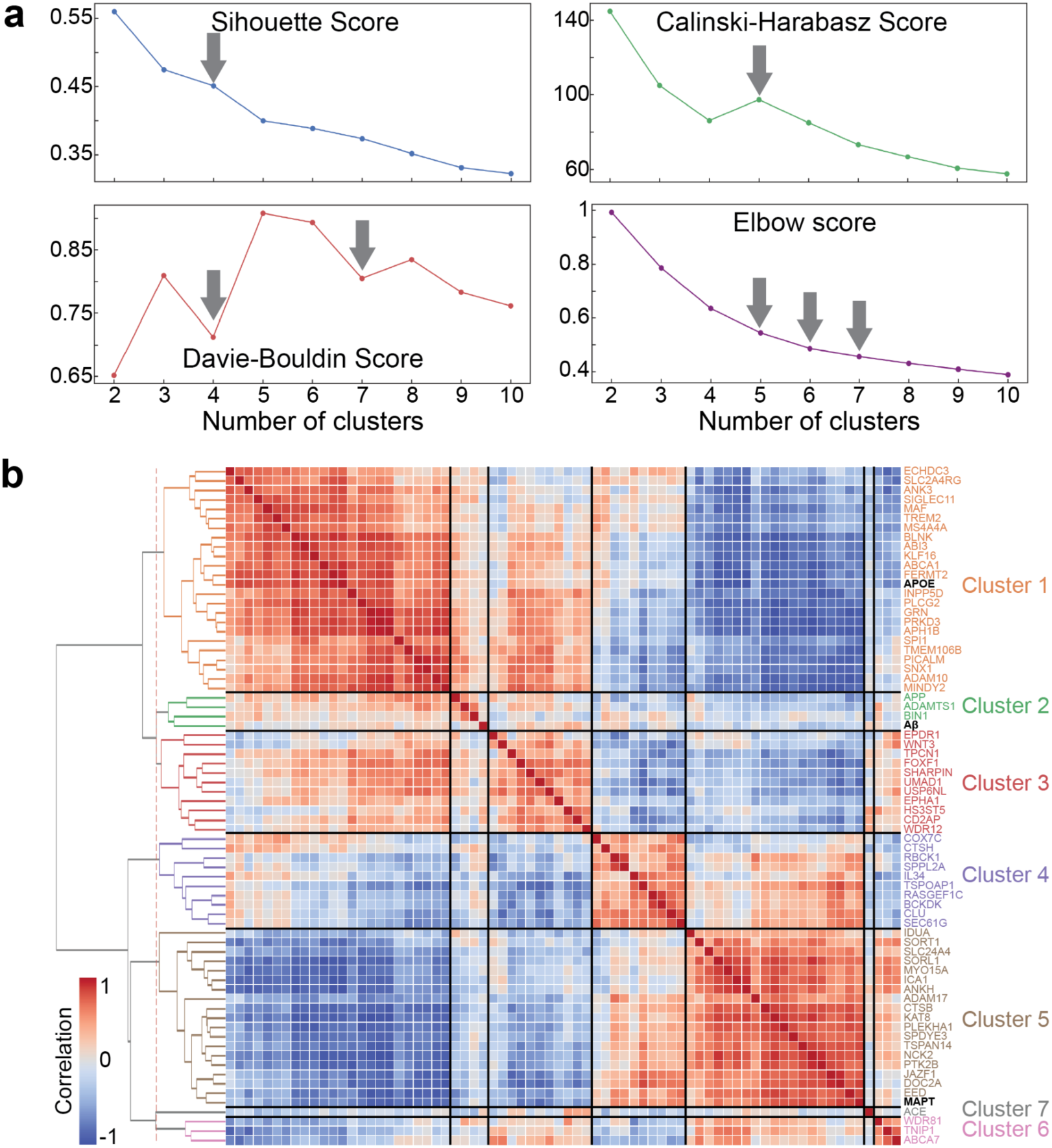
Clusters of AD-related genes identified from GWAS studies. Related to Extended Data Fig. 4. **a)** Four evaluation metrics were used to determine the optimal number of clusters: Silhouette Score (higher values indicate better cohesion and separation), Calinski-Harabasz Index (higher values reflect a better ratio of between-cluster to within-cluster dispersion), Davies-Bouldin Index (lower values indicate less similarity between clusters), and Elbow Score (measures compactness, used in the Elbow Method). The optimal number of clusters for each metric is marked with grey arrows. **b)** Correlation map of the identified genes, with text color-coded by cluster. The three conventional factors are shown in bold black.

**Extended Data Fig. 8.**
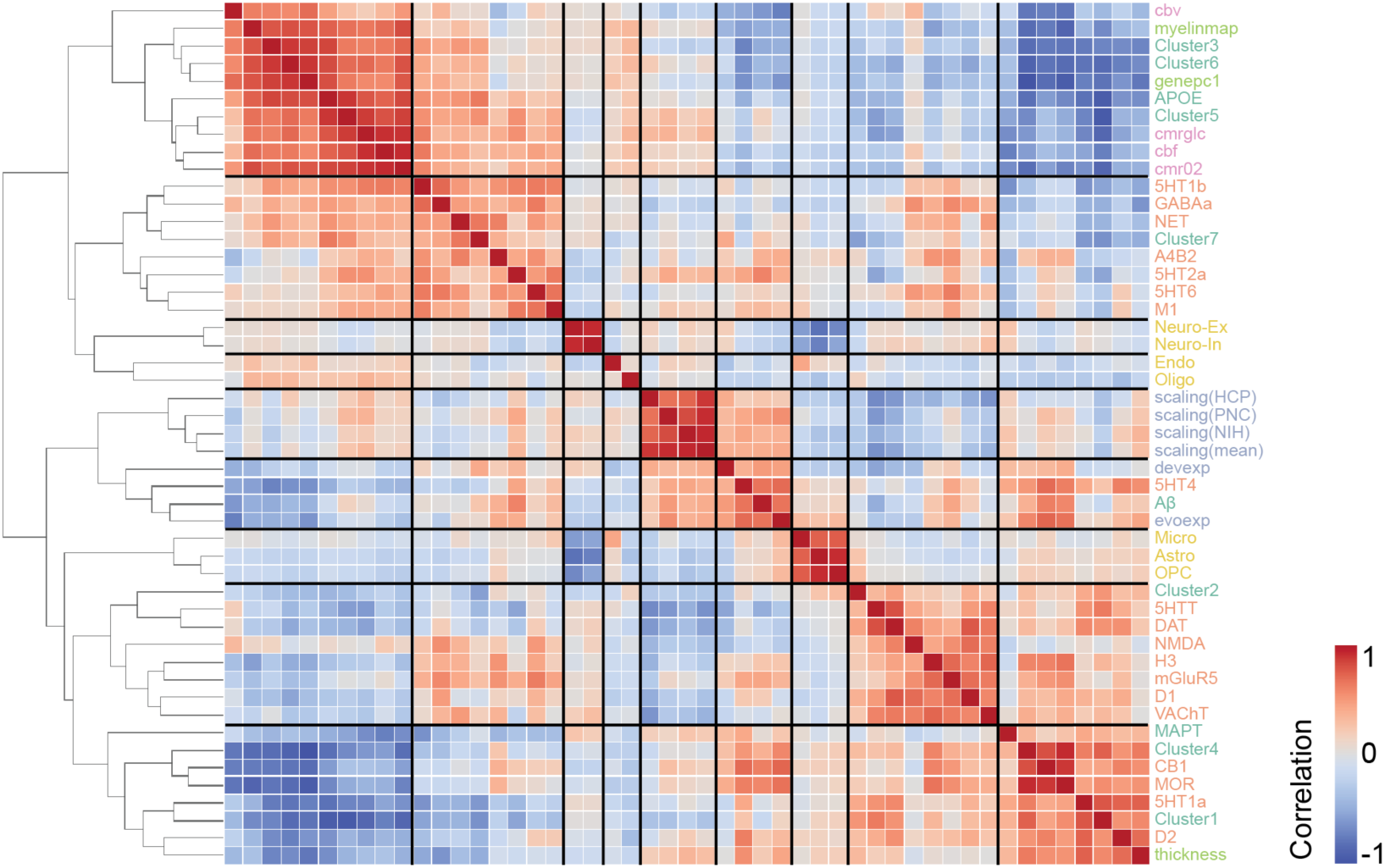
Correlation among 49 regional biological factors. Related to Extended Data Fig. 4. A correlation heatmap matrix with hierarchical clustering reveals relationships among 49 regional biological factors. Nine distinct clusters of factor distributions were identified, with hierarchical trees indicating their groupings. Factors are color-coded by type to highlight their categorization and relationships within clusters. astro = astrocytes; endo = endothelial cells; micro = microglia; neuron-ex = excitatory neurons; neuron-in = inhibitory neurons; oligo = oligodendrocytes; and opc = oligodendrocyte precursor cells.

## Supplementary information

### Supplementary Equation

#### The Susceptible-Infectious-Recovered Model

The model simulates four key processes governing tau dynamics: **spread, synthesis, clearance, and misfolding**. Equations (1)–(4) formally describe how region-specific vulnerability input parameters independently modulate each process, determining the resulting concentrations of normal (*N_i_*) and misfolded (*M_i_*) tau in each region *i* over discrete time steps.

At each simulation step, tau dynamics are updated sequentially through spreading, synthesis, clearance, and misfolding. Protein flux between axonal compartments and regional cell bodies is explicitly modeled during the spreading step, whereas synthesis and clearance operate locally within each region. All state variables at time *t* depend on the protein concentrations from the previous time step (*t* − 1).

During the **spread** process, the normal or misfolded protein concentration in region *i* (*P_i_*) is determined by the proteins retained from the previous time step (*P_i,t_*_−1_) and the flux of proteins exchanged between the axon/terminal (edge) and the soma (region). Specifically, *P_i, incoming_* represents the number of tau proteins transported from the connected axons (presynaptic regions) to the soma of region *i* (postsynaptic target), while *P_i, outgoing_* denotes the number of tau proteins conveyed from the soma toward its axons/terminals within region *i*, reflecting anterograde axonal transport. The extended input parameter **spread_rate***_i_* allows regional modulation of tau spread, controlling the amount of tau transferred from presynaptic regions to the soma of region *i*, scaled by the time increment Δ*t*.

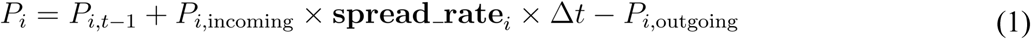

The **synthesis** process follows spreading. The number of newly produced normal proteins in region *i* (*N_i,sythesized_*) is determined by the global synthesis rate across the brain (synthesis_rate*_global_*) and the input region-specific synthesis rate (**synthesis_rate***_i_*). The number of proteins synthesized within a time interval Δ*t* is constant for each region and independent of the existing protein concentration.

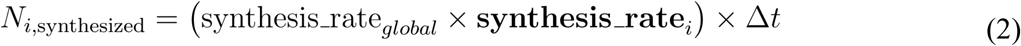

The **clearance** process occurs concurrently with synthesis over a short time interval Δ*t*. The number of normal or misfolded proteins cleared from region *i* (*P_i,cleared_*) depends on the current protein number after spreading (*P_i_*) and the input regional clearance rate (**clearance_rate***_i_*).

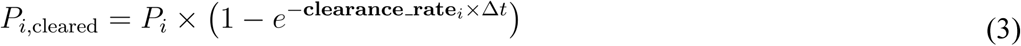

During the **misfolding** process, the number of newly misfolded proteins in region *i* (*M_i,misfolded_*) is determined by the remaining pool of normal proteins after the clearance process (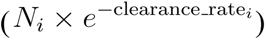), the current level of misfolded protein after the spreading process (*M_i_*), the global misfolding probability (misfold_rate*_global_*), and the input regional misfolding rate (**misfold_rate***_i_*). The extended parameter **misfold_rate***_i_* provides regional control over the misfolding propensity of tau.

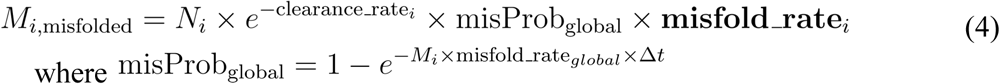

During each simulation step, the combined effects of these processes determine the updated levels of normal (*N_i_*) and misfolded (*M_i_*) proteins in region *i* at time *t*. The resulting concentrations are expressed as:

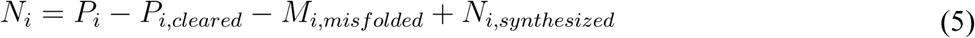

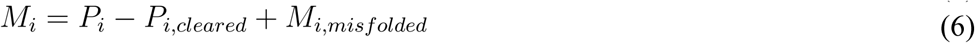

### Supplementary Notes

#### Supplementary Note 1. Participant inclusion and diagnostic criteria

The Swedish BioFINDER-2 study enrolls participants across multiple clinical stages of cognitive impairment. The present study included participants from Cohorts A-D; individuals with non–Alzheimer’s disease neurodegenerative disorders (Cohort E) were not included.

Cohorts A and B comprised neurologically and cognitively unimpaired individuals aged 40–65 years (Cohort A) or 66–100 years (Cohort B). Inclusion criteria were: (i) absence of cognitive symptoms as assessed by a physician with expertise in cognitive disorders; (ii) Mini-Mental State Examination (MMSE) score of 27–30 (Cohort A) or 26–30 (Cohort B) at screening; (iii) no fulfillment of criteria for mild cognitive impairment (MCI) or dementia according to DSM-5; and (iv) fluency in Swedish. Recruitment of these cohorts was designed to achieve approximately equal proportions of APOE ε4 carriers.

Cohort C included participants aged 40–100 years referred to memory clinics due to cognitive complaints. Inclusion criteria were: (i) MMSE score of 24–30; (ii) absence of dementia according to DSM-5; and (iii) fluency in Swedish. Participants were classified as having MCI if they performed worse than −1.5 standard deviations relative to age- and education-adjusted norms in at least one cognitive domain. Cognitive assessment covered attention/executive function (Trail Making Test A and B, Symbol Digit Modalities Test, AQT), memory (immediate and delayed recall from the Alzheimer’s Disease Assessment Scale), language (verbal fluency and short Boston Naming Test), and visuospatial function (Visual Object and Space Perception battery). Participants who did not meet MCI criteria were classified as having subjective cognitive decline (SCD). In accordance with the NIA–AA research framework, individuals with SCD were grouped together with cognitively unimpaired participants in analyses where appropriate.

Cohort D comprised participants aged 40–100 years referred to memory clinics who fulfilled DSM-5 criteria for dementia (major neurocognitive disorder) due to Alzheimer’s disease. Inclusion criteria included an MMSE score ≥12 and fluency in Swedish.

Across all cohorts, exclusion criteria were: (i) significant unstable systemic illness interfering with study participation; (ii) current significant alcohol or substance misuse; and (iii) refusal to undergo lumbar puncture, MRI, or PET imaging.

#### Supplementary Note 2. Details of PET acquisition and preprocessing

Tau PET imaging was performed using [^18^F]RO948 on a digital scanner (Discovery MI; GE Healthcare). PET data were acquired in 70 to 90 minutes post-injection.

Low-dose computed tomography scans were acquired immediately prior to PET imaging for attenuation correction. Image reconstruction was performed using VPFX-S (ordered subset expectation maximization combined with corrections for time-of-flight and point spread function) with 6 iterations, 17 subsets, and applied 3-mm Gaussian smoothing, a standard Z filter, and a 25.6-cm field of view (256 × 256 matrix). List-mode data were binned into four consecutive 5-minute frames.

PET frames were motion corrected using rigid-body realignment implemented in AFNI (*3dvolreg*), summed within participant, and rigidly coregistered to the corresponding native-space T1-weighted MRI. Subsequent intensity normalization using an inferior cerebellar gray matter reference region yielded standardized uptake value ratio (SUVR) images, which were used for regional quantification as described in the main Methods.

#### Supplementary Note 3. Hyperparameter tuning

For all SIR model simulations, hyperparameters were systematically tuned to optimize correspondence between simulated and observed tau patterns. The parameter search space included the probability of tau remaining within the neuronal cell body (**stay probability**; values: 0.01, 0.05, 0.1, 0.3, 0.5, 0.7, 0.9), the global tau misfolding rate (**transform rate**; values: 0.1, 0.5, 0.9, 1.0, 1.5, 1.9, 2.0, 2.5, 2.9, 3.0), and tau spreading velocity along anatomical connections (**velocity**; values: 0.1, 0.3, 0.5, 0.7, 0.9, 1.0). For each combination of parameters, model performance was evaluated based on its ability to reconstruct group-averaged regional tau presence (TPP) or tau load (SUVR). The parameter set yielding the best fit was then selected for subsequent analyses.

#### Supplementary Note 4. Epidemic spreading model

The spread of tau pathology across connected brain regions was simulated using the Epidemic Spreading Model (ESM), a previously described diffusion-based framework applied to misfolded protein propagation in neurodegenerative disease. The ESM models the diffusion of a pathological signal from a predefined epicenter through a weighted brain connectome over time. Model dynamics are governed by subject-specific parameters describing global tau production rate, clearance rate, and estimated age of onset, which modulate the overall extent of spread without determining regional patterning.

The ESM was fit within-subject using regional tau presence (TPP) or tau load (SUVR) as input, represented as region-by-subject matrices. For the primary analyses, the left and right entorhinal cortex were selected as model epicenters based on neuropathological evidence of early tau accumulation. Model predictions were compared to observed regional tau patterns, and performance was quantified based on the correspondence between predicted and observed values. Consistent with prior work, the free parameters primarily controlled global tau burden, allowing the model to accommodate individuals across the Alzheimer’s disease spectrum.

#### Supplementary Note 5. *Franzmeier et al., 2020* statistical approach

To further assess the relationship between anatomical connectivity and tau propagation, we performed connectivity-based statistical analyses following previously described approaches. Cross-sectionally estimated tau spreading matrices were constructed by concatenating regional tau presence or tau load across Aβ-positive participants. Rows (subjects) and columns (regions) were rank-ordered by their respective means, yielding tau positivity sequences.

Epicenters were defined as the 10% of regions exhibiting the earliest tau involvement. Mean connectivity-based distance from these epicenters was calculated using the structural connectome employed in the SIR and ESM analyses. Associations between tau spreading sequences and connectivity-based distances were assessed using linear regression.

### Supplementary Figures

**Fig. S1.**
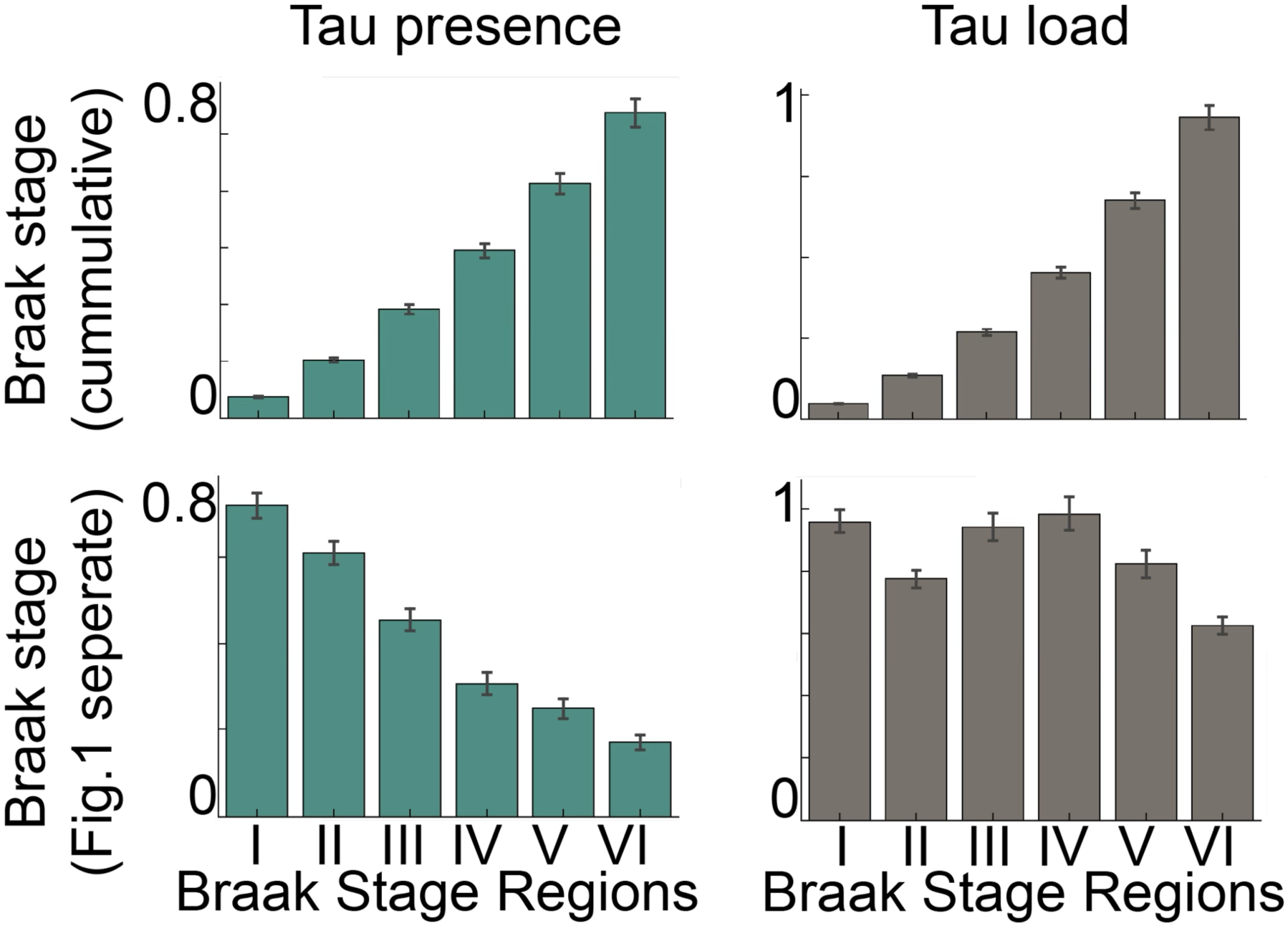
Accumulative versus separate Braak stage definitions for tau presence and load. Mean ± SEM values of tau presence (left) and tau load (right) across Braak stage regions. Top: values calculated accumulatively, including regions from earlier stages. Bottom: values calculated separately, including only regions of the corresponding stage (approach used in the main manuscript, Fig. 1). The latter approach is a better approximation to Braak staging, where braak stage regions are addressed independently.

**Fig. S2.**
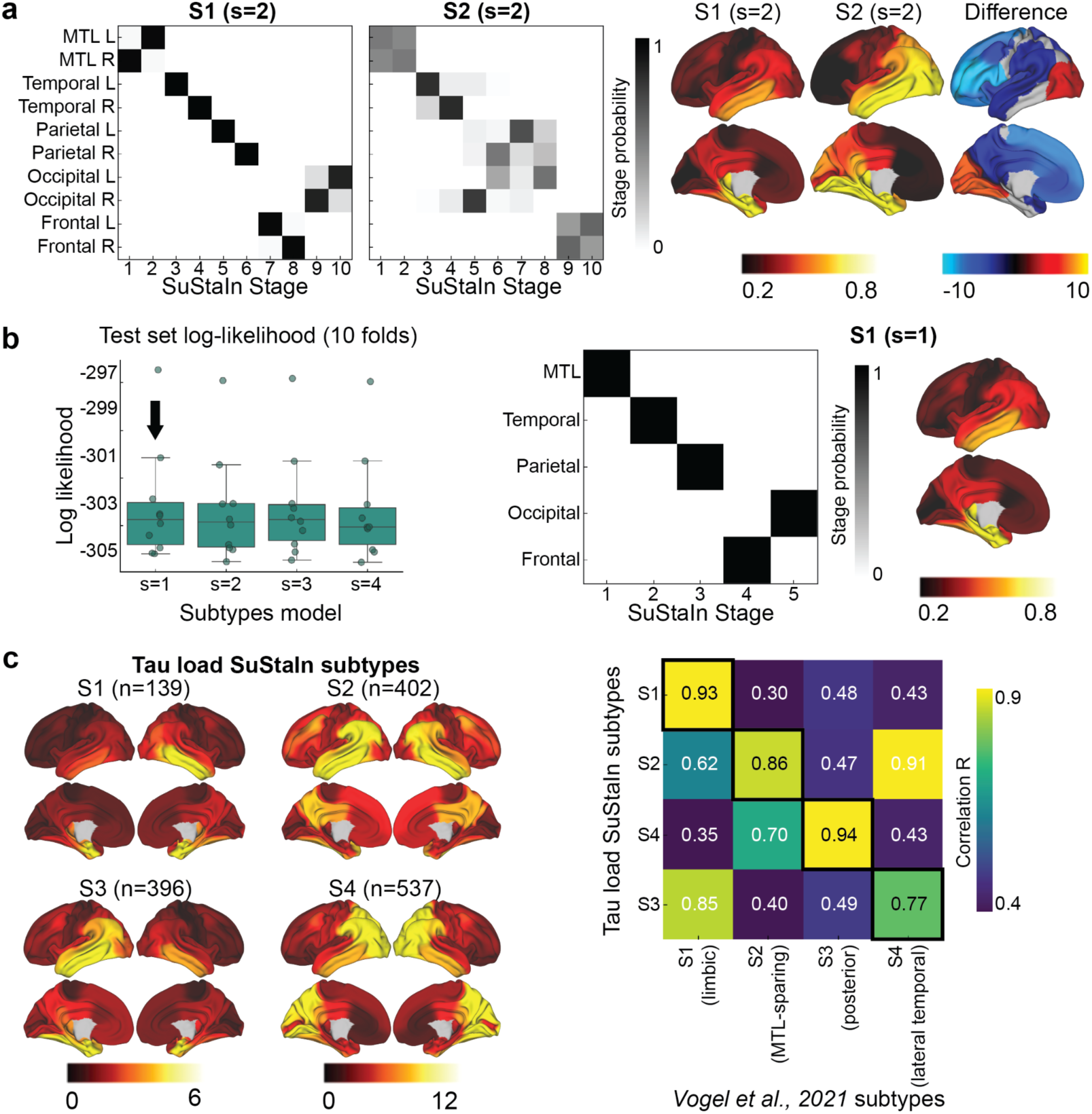
Tau presence is consistent across individuals, tau load is heterogeneous, Related to main text Fig. 1 and Fig. 5. **a)** Left: Positional variance for two identified tau presence subtypes using 10 brain regions. Each box represents the confidence that a brain region has reached a certain pathology level at a given SuStaIn (disease progression) stage, with darker colors indicating higher certainty. Right: Tau presence cortical surface projections for the two subtypes and a difference map. **b)** Results using five brain regions. Left: For each subtype model (k=1–4), the distribution of average negative log-likelihood across cross-validation folds of left-out individuals is shown (higher log-likelihood indicates better model fit). Middle: Positional variance for two tau presence subtypes. Right: Tau presence cortical surface projections of the identified tau presence subtype. **c)** Right: Brain map for the identified four tau load subtypes. Left: Confusion matrix comparing subtypes identified in the current study (SuStaIn subtypes; y-axis) with those from *Vogel et al., 2021* (x-axis). Values represent spatial correlations between average regional tau for each subtype. Diagonal values indicate subtype similarity across both parameter sets. L: left; R: right; SuStaIn: Subtype and Stage Inference model.

**Fig. S3.**
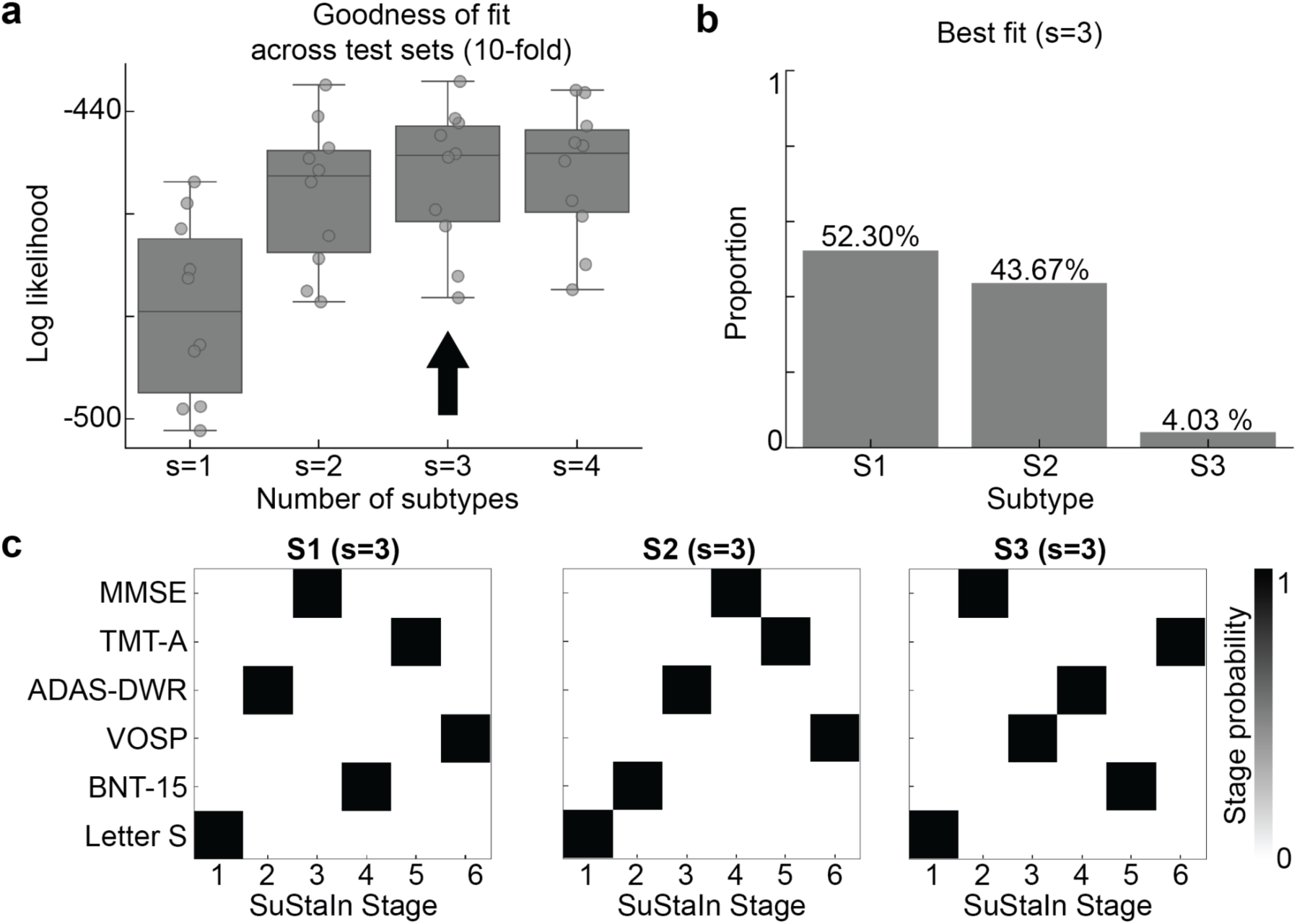
SuStaIn-derived three spatiotemporal patterns for cognitive positive-probabilty. Related to main text. **Fig. 1**. To prove that the binarizing data does not automatically lead to a lack of subtypes, binarized cognitive data and ran SuStaIn on this data. Six cognitive measurements were first transformed into cognitive positive-probability (presence of cognitive domain impairment) using two-component Gaussian Mixture model, following the same exact procedure performed to calculate tau presence. A SuStaIn model was then applied on binarized cognitive impairment of six measures. **a)** For each subtype model (s=1-4), distributions of average negative log-likelihood across 10 cross-validation folds of left-out individuals are shown, higher values indicate better fit. The best-fit model is marked by the black arrow. **b)** Bar plots show the proportion of participants assigned to each subtype. **c)** Positional variance for three cognitive domain impairment sequence subtypes. SuStaIn: Subtype and Stage Inference; MMSE: mini-mental state examination; TMT-A: trail making test A; ADAS-DWR: Alzheimer’s Disease Assessment Scale, delayed word recall; VOSP: Visual Object and Space Perception Battery; BNT-15: Boston Naming Test - short form; Letter S: letter S fluency test.

**Fig. S4.**
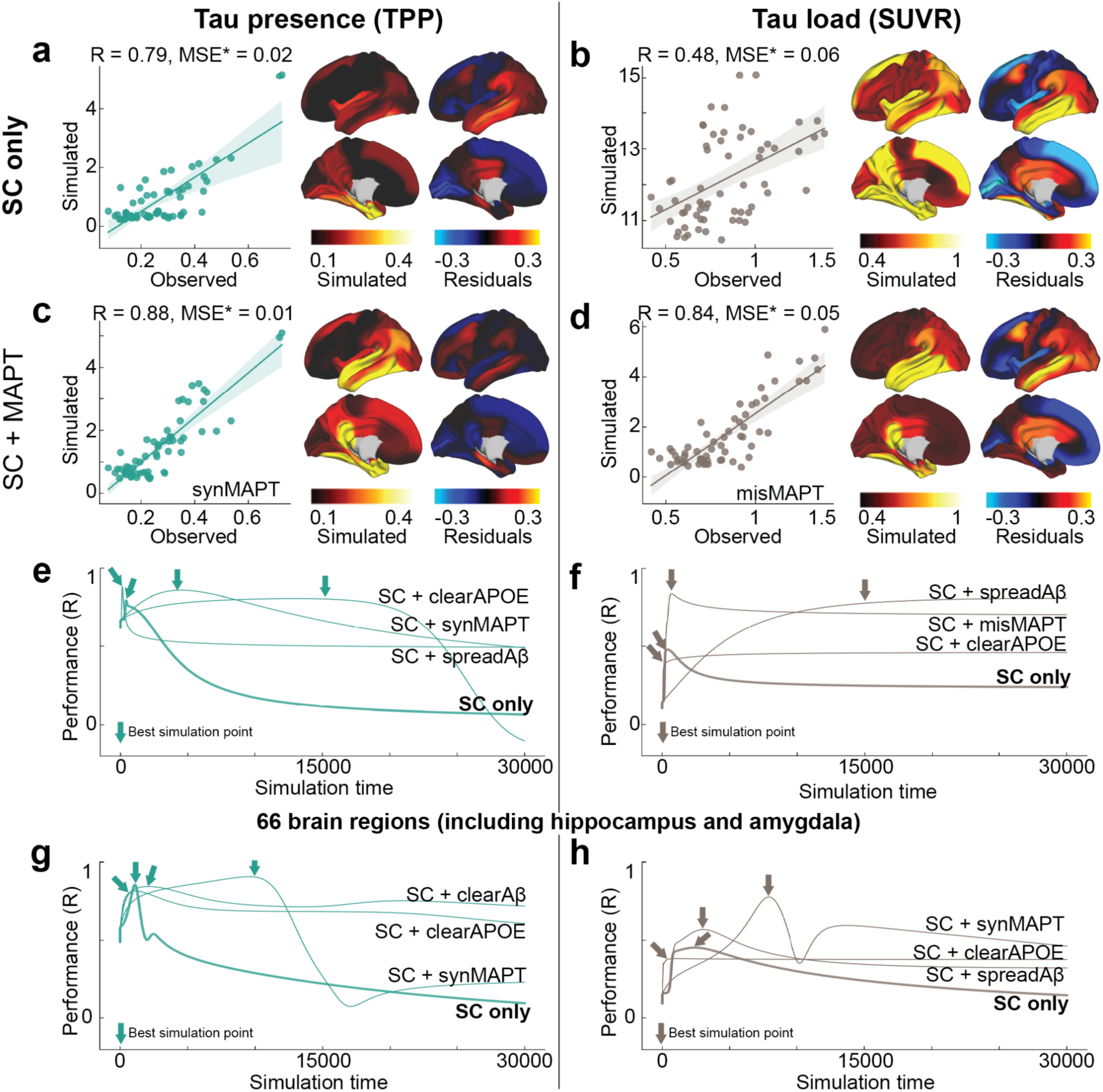
Replication of main text Fig. 2 using a different structural connectome. Results of the SIR model simulating tau diffusion through a structural connectome (derived from healthy young individuals from HCP) using an entorhinal cortex epicenter. Note that hippocampus and amygdala are not included in this connectome. **a, b)** Model performance for tau presence (TPP) a), and tau load (SUVR) b). Scatter plots compare SIR model predictions to observed tau TPP/SUVR, while cortical surface projections show simulated tau presence and model residuals. Residuals represent the difference between observed tau values and simulated tau values, with the simulated values normalized to the range of observed values. **c-h)** To assess the influence of regional vulnerability on tau propagation, three regional properties were incorporated into the model: *MAPT* expression, *APOE* expression, and A*β* deposition. c, d) The best-fitting regional property influencing tau presence (SC plus *MAPT* expression affecting regional synthesis) is shown in c), and for tau load in d). **e, f)** Model performance (SC only vs. SC with three regional properties) over simulated time for TPP and SUVR, respectively. **g, h)** Model performance (SC only vs. SC with three regional properties, simulated with 66 brain regions used in Fig. 2) over simulated time for TPP and SUVR, respectively. MSE* was calculated using normalized simulated values (scaled to the range of observed tau values) to ensure a fair comparison, as the model lacks constraints on simulated values, which can increase without limit. SUVR: standardized uptake value ratio; TPP: tau positive-probability; SC: structural connectivity; R: Pearson correlation coefficient; MSE: mean squared error; *MAPT*: microtubule associated protein tau; *APOE*: Apolipoprotein E; syn: synthesis; clear: clearance; mis: misfold.

### Supplementary Tables

**Table S1.** Sample characteristics.

|  | **BioFINDER2** |
| --- | --- |
| **Participants (N)** | 646 |
| **Diagnosis**<br>**(%Normal/MCI/AD)** | 33.90/32.82/33.28 |
| **Age (yrs)** | 73.27 (7.73) [44.26 - 93.26] |
| **Gender (F/M)** | 325/321 |
| **Education (yrs)** | 12.61 (4.12) [3.00 - 33.00] |
| **APOE4 (carrier/non-carrier)** | 408/178 |
| **CSF Aβ42/40** | 0.05 (0.01) [0.02 - 0.08] |
| **CSF p-tau217 (pg/ml)** | 403.24 (327.35) [19.10 - 2016.15] |

Mean (standard deviation) [minimum, maximum] values are reported for continuous variables, and participant counts are provided for categorical variables (Participants, Diagnosis, Gender, APOE4).
% = percentage; MCI = Mild Cognitive Impairment; AD = Alzheimer's Disease; yrs = years old; F = female; M = male; CSF = cerebral spinal fluid; Aβ + = $\beta$ amyloid positive

**Table S2.** Model performance for simulating tau progression along anatomical connections (66 regions, including hippocampas and amygdala), influenced by regional classical AD-related elements. Related to Fig. 2.

| Models | Tau presence (TPP) |  |  |  | Tau load (SUVR) |  |  |  |
| --- | --- | --- | --- | --- | --- | --- | --- | --- |
|  | R | MSE* | Hyperparams | Time | R | MSE* | Hyperparams | Time |
| **SC only** | **0.8560** | **0.0195** | (0.7, 1.9, 0.1) | 1101 | **0.4534** | **0.0723** | (0.9, 1.5, 0.1) | 2412 |
| +spreadMAPT | 0.6843 | 0.0310 | (0.9, 1.5, 0.1) | 742 | 0.4803 | **0.0604** | (0.9, 1.9, 0.1) | 4372 |
| +synMAPT | **0.9083** | **0.0102** | (0.05, 2.5, 0.1) | 9540 | **0.7777** | 0.0615 | (0.01, 3, 0.1) | 7898 |
| +misMAPT | 0.7495 | 0.0252 | (0.1, 3, 0.5) | 2044 | 0.7017 | 0.0659 | (0.3, 2.9, 0.5) | 1836 |
| +clearMAPT | 0.8404 | 0.0233 | (0.1, 0.5, 0.9) | 1970 | 0.2848 | 0.1743 | (0.9, 0.1, 0.1) | 248 |
| +spreadAβ | 0.6833 | **0.0268** | (0.9, 1.5, 0.1) | 765 | **0.5694** | **0.0546** | (0.9, 1.5, 0.1) | 3048 |
| +synAβ | 0.7347 | 0.0358 | (0.01, 2, 0.1) | 11757 | 0.4280 | 0.1506 | (0.9, 3, 1) | 69 |
| +clearAβ | 0.7235 | 0.0312 | (0.01, 3, 0.1) | 9694 | 0.4590 | 0.1498 | (0.5, 2.9, 0.1) | 2943 |
| +misAβ | **0.8447** | 0.0240 | (0.05, 0.5, 0.7) | 2101 | 0.2894 | 0.1605 | (0.01, 0.5, 0.1) | 16812 |
| +spreadAPOE | 0.7475 | 0.0262 | (0.9, 0.1, 1) | 29999 | 0.3384 | **0.0737** | (0.9, 1.5, 0.1) | 1077 |
| +synAPOE | 0.7171 | 0.0282 | (0.01, 1, 1) | 4512 | 0.3296 | 0.0865 | (0.01, 3, 0.7) | 29999 |
| +misAPOE | 0.7140 | 0.0370 | (0.01, 1.5, 0.1) | 13202 | 0.2558 | 0.1713 | (0.01, 1, 0.1) | 16248 |
| +clearAPOE | **0.8161** | **0.0223** | (0.05, 0.5, 1) | 1158 | **0.3829** | 0.1018 | (0.9, 3, 1) | 1112 |

Each regional canonical AD-related element was parameterized to influence tau simulation via spread, synthesis, misfolding, and clearance rates. All models underwent hyperparameter tuning, and the best-performing model was selected based on the highest simulation accuracy at the optimal time point (total time = 30,000) and hyperparameter set. The best-performing model for each AD-related element is highlighted in bold. Selected hyperparameters, including stay probability, transform rate, and velocity, are shown.
TPP = tau-positive probability; SUVR = standardized uptake ratio; R = Pearson correlation coefficient; MSE = mean squared error; SC = structural connectivity; syn = synthesis; mis = misfold; clear = clearance; Aβ = $\beta$ amyloid

**Table S3.**
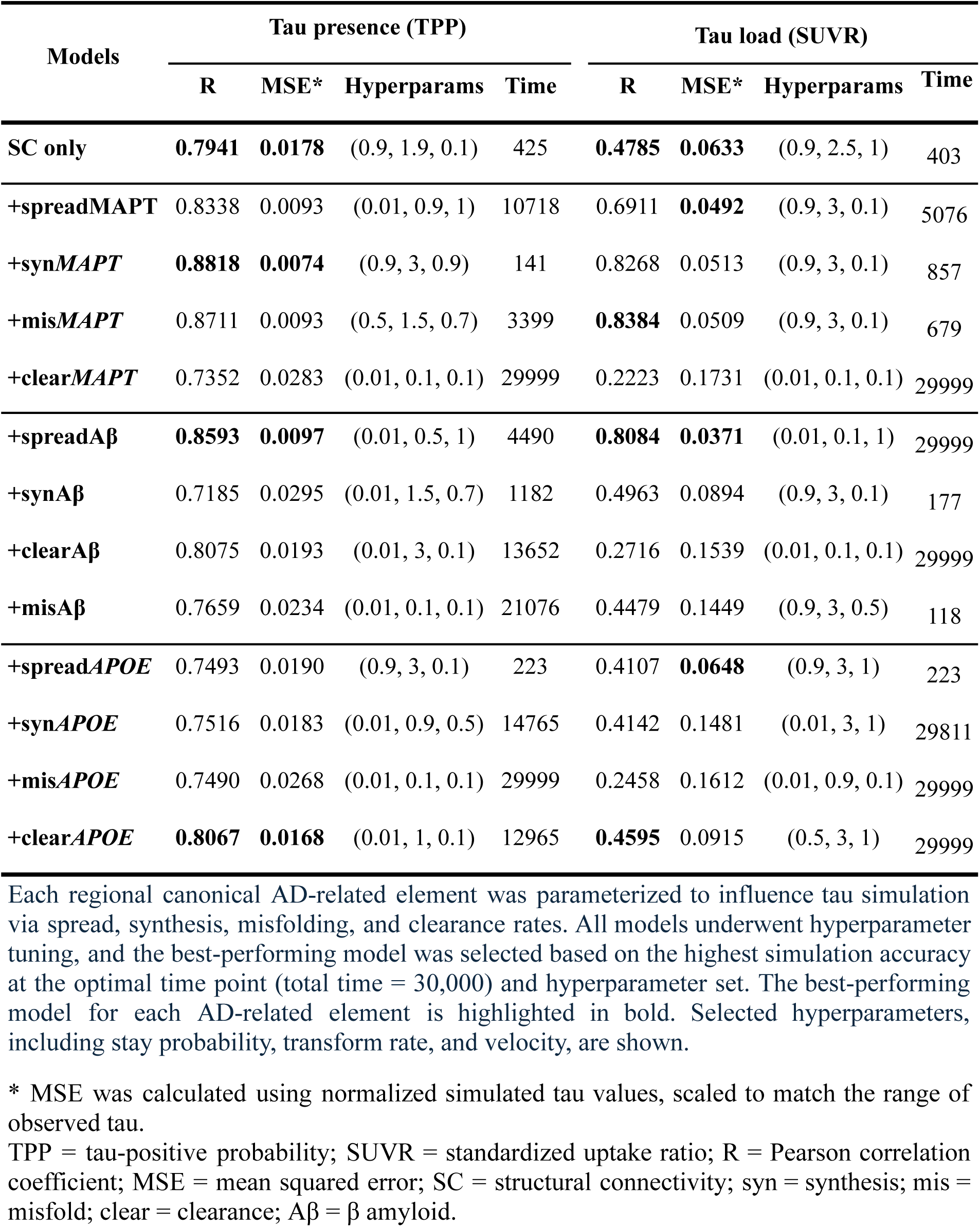
Model performance for simulating tau progression along anatomical connections (62 regions, excluding hippocampas and amygdala), influenced by regional classical AD-related elements. Related to Fig. S4.

| Models | Tau presence (TPP) |  |  |  | Tau load (SUVR) |  |  |  |
| --- | --- | --- | --- | --- | --- | --- | --- | --- |
|  | R | MSE* | Hyperparams | Time | R | MSE* | Hyperparams | Time |
| **SC only** | **0.7941** | **0.0178** | (0.9, 1.9, 0.1) | 425 | **0.4785** | **0.0633** | (0.9, 2.5, 1) | 403 |
| +spreadMAPT | 0.8338 | 0.0093 | (0.01, 0.9, 1) | 10718 | 0.6911 | **0.0492** | (0.9, 3, 0.1) | 5076 |
| +synMAPT | **0.8818** | **0.0074** | (0.9, 3, 0.9) | 141 | 0.8268 | 0.0513 | (0.9, 3, 0.1) | 857 |
| +misMAPT | 0.8711 | 0.0093 | (0.5, 1.5, 0.7) | 3399 | **0.8384** | 0.0509 | (0.9, 3, 0.1) | 679 |
| +clearMAPT | 0.7352 | 0.0283 | (0.01, 0.1, 0.1) | 29999 | 0.2223 | 0.1731 | (0.01, 0.1, 0.1) | 29999 |
| +spreadAβ | **0.8593** | **0.0097** | (0.01, 0.5, 1) | 4490 | **0.8084** | **0.0371** | (0.01, 0.1, 1) | 29999 |
| +synAβ | 0.7185 | 0.0295 | (0.01, 1.5, 0.7) | 1182 | 0.4963 | 0.0894 | (0.9, 3, 0.1) | 177 |
| +clearAβ | 0.8075 | 0.0193 | (0.01, 3, 0.1) | 13652 | 0.2716 | 0.1539 | (0.01, 0.1, 0.1) | 29999 |
| +misAβ | 0.7659 | 0.0234 | (0.01, 0.1, 0.1) | 21076 | 0.4479 | 0.1449 | (0.9, 3, 0.5) | 118 |
| +spreadAPOE | 0.7493 | 0.0190 | (0.9, 3, 0.1) | 223 | 0.4107 | **0.0648** | (0.9, 3, 1) | 223 |
| +synAPOE | 0.7516 | 0.0183 | (0.01, 0.9, 0.5) | 14765 | 0.4142 | 0.1481 | (0.01, 3, 1) | 29811 |
| +misAPOE | 0.7490 | 0.0268 | (0.01, 0.1, 0.1) | 29999 | 0.2458 | 0.1612 | (0.01, 0.9, 0.1) | 29999 |
| +clearAPOE | **0.8067** | **0.0168** | (0.01, 1, 0.1) | 12965 | **0.4595** | 0.0915 | (0.5, 3, 1) | 29999 |

Each regional canonical AD-related element was parameterized to influence tau simulation via spread, synthesis, misfolding, and clearance rates. All models underwent hyperparameter tuning, and the best-performing model was selected based on the highest simulation accuracy at the optimal time point (total time = 30,000) and hyperparameter set. The best-performing model for each AD-related element is highlighted in bold. Selected hyperparameters, including stay probability, transform rate, and velocity, are shown.
TPP = tau-positive probability; SUVR = standardized uptake ratio; R = Pearson correlation coefficient; MSE = mean squared error; SC = structural connectivity; syn = synthesis; mis = misfold; clear = clearance; Aβ = $\beta$ amyloid.

**Table S4. Performance of models better than null for tau presence and load, Related to Fig. 3**.

**(Attached Supplementary_Table4_Presence-Load_mdoels_performance.xlsx)**

The optimal time points (total time = 30,000) are provided, along with the selected hyperparameters: stay probability, transform rate, and velocity.

*MSE was calculated using normalized simulated tau values, scaled to match the range of observed tau.

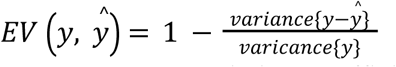

R = Pearson correlation coefficient; EV = explained variance; MSE = mean squared error; syn = synthesis; mis = misfold; clear = clearance; Aβ = β amyloid; SC = structural connectivity; FC = functional connectivity; GC = gene coexpression; NS = neurotransmission markerreceptor similarity; LS = laminar similarity; MC = metabolic covariance; EP = electrophysiological; TS = temporal similarity; CBF: cerebral blood flow; CBV: cerebral blood volume; thickness: cortical thickness; scaling: allometric scalling.

**Table S5.**
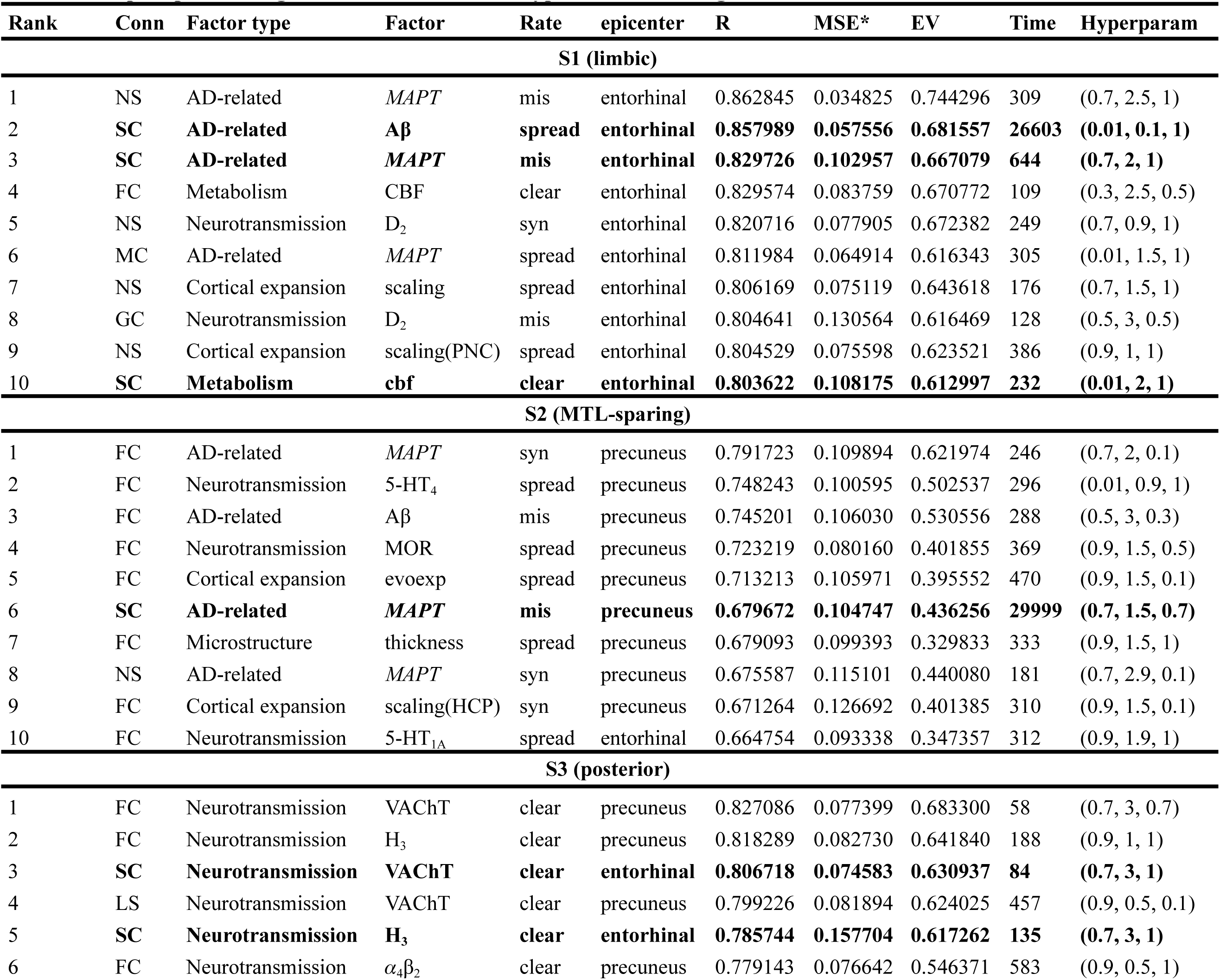
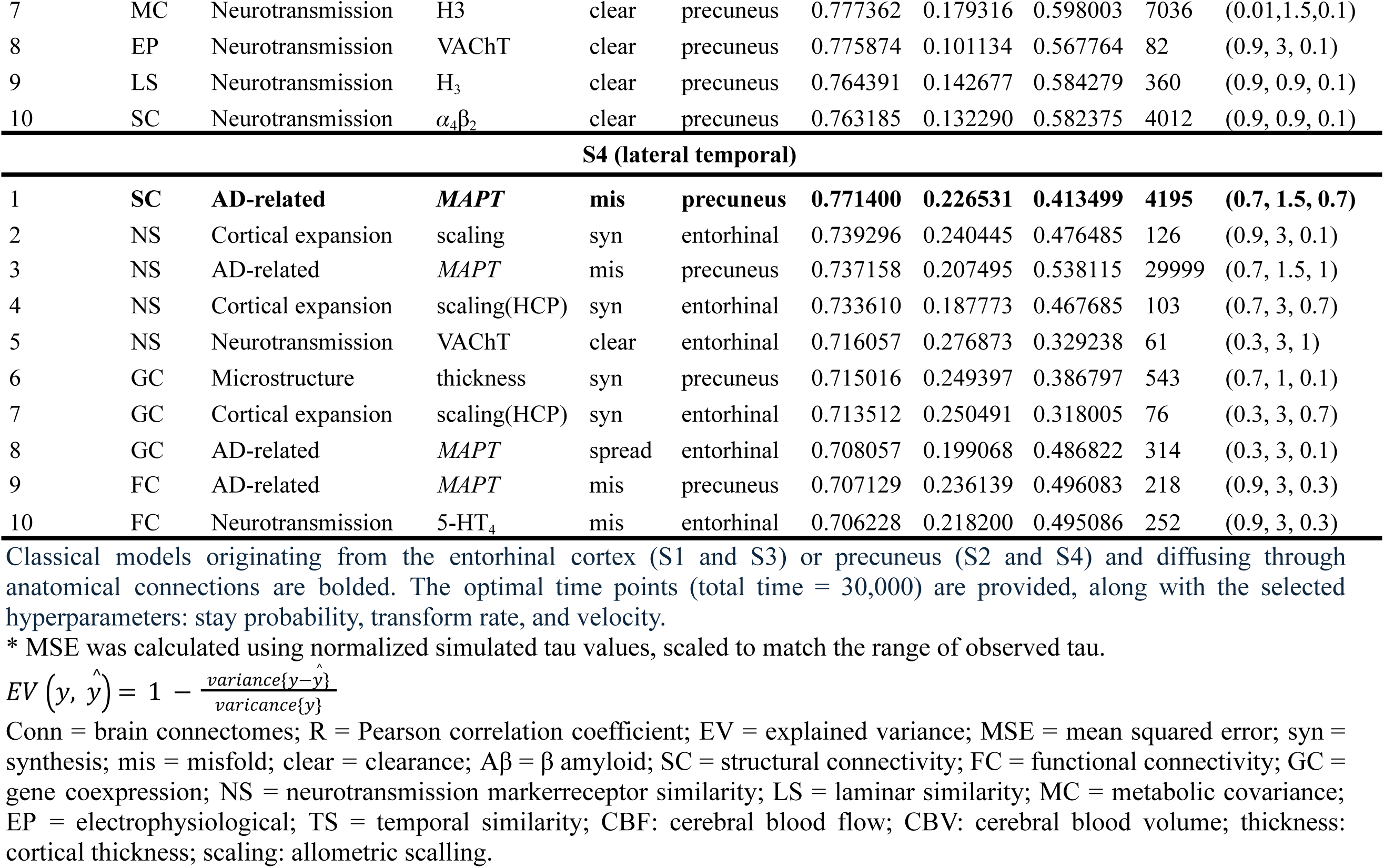
Top 10 performing models for four tau subtypes. Related to Fig. 5.

| Rank | Conn | Factor type | Factor | Rate | epicenter | R | MSE* | EV | Time | Hyperparam |
| --- | --- | --- | --- | --- | --- | --- | --- | --- | --- | --- |
| **S1 (limbic)** |  |  |  |  |  |  |  |  |  |  |
| 1 | NS | AD-related | *MAPT* | mis | entorhinal | 0.862845 | 0.034825 | 0.744296 | 309 | (0.7, 2.5, 1) |
| 2 | **SC** | **AD-related** | **Aβ** | **spread** | **entorhinal** | **0.857989** | **0.057556** | **0.681557** | **26603** | **(0.01, 0.1, 1)** |
| 3 | **SC** | **AD-related** | ***MAPT*** | **mis** | **entorhinal** | **0.829726** | **0.102957** | **0.667079** | **644** | **(0.7, 2, 1)** |
| 4 | FC | Metabolism | CBF | clear | entorhinal | 0.829574 | 0.083759 | 0.670772 | 109 | (0.3, 2.5, 0.5) |
| 5 | NS | Neurotransmission | D<sub>2</sub> | syn | entorhinal | 0.820716 | 0.077905 | 0.672382 | 249 | (0.7, 0.9, 1) |
| 6 | MC | AD-related | *MAPT* | spread | entorhinal | 0.811984 | 0.064914 | 0.616343 | 305 | (0.01, 1.5, 1) |
| 7 | NS | Cortical expansion | scaling | spread | entorhinal | 0.806169 | 0.075119 | 0.643618 | 176 | (0.7, 1.5, 1) |
| 8 | GC | Neurotransmission | D<sub>2</sub> | mis | entorhinal | 0.804641 | 0.130564 | 0.616469 | 128 | (0.5, 3, 0.5) |
| 9 | NS | Cortical expansion | scaling(PNC) | spread | entorhinal | 0.804529 | 0.075598 | 0.623521 | 386 | (0.9, 1, 1) |
| 10 | **SC** | **Metabolism** | **cbf** | **clear** | **entorhinal** | **0.803622** | **0.108175** | **0.612997** | **232** | **(0.01, 2, 1)** |
| **S2 (MTL-sparing)** |  |  |  |  |  |  |  |  |  |  |
| 1 | FC | AD-related | *MAPT* | syn | precuneus | 0.791723 | 0.109894 | 0.621974 | 246 | (0.7, 2, 0.1) |
| 2 | FC | Neurotransmission | 5-HT<sub>4</sub> | spread | precuneus | 0.748243 | 0.100595 | 0.502537 | 296 | (0.01, 0.9, 1) |
| 3 | FC | AD-related | Aβ | mis | precuneus | 0.745201 | 0.106030 | 0.530556 | 288 | (0.5, 3, 0.3) |
| 4 | FC | Neurotransmission | MOR | spread | precuneus | 0.723219 | 0.080160 | 0.401855 | 369 | (0.9, 1.5, 0.5) |
| 5 | FC | Cortical expansion | evoexp | spread | precuneus | 0.713213 | 0.105971 | 0.395552 | 470 | (0.9, 1.5, 0.1) |
| 6 | **SC** | **AD-related** | ***MAPT*** | **mis** | **precuneus** | **0.679672** | **0.104747** | **0.436256** | **29999** | **(0.7, 1.5, 0.7)** |
| 7 | FC | Microstructure | thickness | spread | precuneus | 0.679093 | 0.099393 | 0.329833 | 333 | (0.9, 1.5, 1) |
| 8 | NS | AD-related | *MAPT* | syn | precuneus | 0.675587 | 0.115101 | 0.440080 | 181 | (0.7, 2.9, 0.1) |
| 9 | FC | Cortical expansion | scaling(HCP) | syn | precuneus | 0.671264 | 0.126692 | 0.401385 | 310 | (0.9, 1.5, 0.1) |
| 10 | FC | Neurotransmission | 5-HT<sub>1A</sub> | spread | entorhinal | 0.664754 | 0.093338 | 0.347357 | 312 | (0.9, 1.9, 1) |
| **S3 (posterior)** |  |  |  |  |  |  |  |  |  |  |
| 1 | FC | Neurotransmission | VAcHT | clear | precuneus | 0.827086 | 0.077399 | 0.683300 | 58 | (0.7, 3, 0.7) |
| 2 | FC | Neurotransmission | H<sub>3</sub> | clear | precuneus | 0.818289 | 0.082730 | 0.641840 | 188 | (0.9, 1, 1) |
| 3 | **SC** | **Neurotransmission** | **VAcHT** | **clear** | **entorhinal** | **0.806718** | **0.074583** | **0.630937** | **84** | **(0.7, 3, 1)** |
| 4 | LS | Neurotransmission | VAcHT | clear | precuneus | 0.799226 | 0.081894 | 0.624025 | 457 | (0.9, 0.5, 0.1) |
| 5 | **SC** | **Neurotransmission** | **H<sub>3</sub>** | **clear** | **entorhinal** | **0.785744** | **0.157704** | **0.617262** | **135** | **(0.7, 3, 1)** |
| 6 | FC | Neurotransmission | $\alpha_4\beta_2$ | clear | precuneus | 0.779143 | 0.076642 | 0.546371 | 583 | (0.9, 0.5, 1) |
| 7 | MC | Neurotransmission | H3 | clear | precuneus | 0.777362 | 0.179316 | 0.598003 | 7036 | (0.01,1.5,0.1) |
| 8 | EP | Neurotransmission | VACHT | clear | precuneus | 0.775874 | 0.101134 | 0.567764 | 82 | (0.9, 3, 0.1) |
| 9 | LS | Neurotransmission | H<sub>3</sub> | clear | precuneus | 0.764391 | 0.142677 | 0.584279 | 360 | (0.9, 0.9, 0.1) |
| 10 | SC | Neurotransmission | $\alpha_4\beta_2$ | clear | precuneus | 0.763185 | 0.132290 | 0.582375 | 4012 | (0.9, 0.9, 0.1) |
| S4 (lateral temporal) |  |  |  |  |  |  |  |  |  |  |
| 1 | SC | AD-related | *MAPT* | mis | **precuneus** | **0.771400** | **0.226531** | **0.413499** | **4195** | **(0.7, 1.5, 0.7)** |
| 2 | NS | Cortical expansion | scaling | syn | entorhinal | 0.739296 | 0.240445 | 0.476485 | 126 | (0.9, 3, 0.1) |
| 3 | NS | AD-related | *MAPT* | mis | precuneus | 0.737158 | 0.207495 | 0.538115 | 29999 | (0.7, 1.5, 1) |
| 4 | NS | Cortical expansion | scaling(HCP) | syn | entorhinal | 0.733610 | 0.187773 | 0.467685 | 103 | (0.7, 3, 0.7) |
| 5 | NS | Neurotransmission | VACHT | clear | entorhinal | 0.716057 | 0.276873 | 0.329238 | 61 | (0.3, 3, 1) |
| 6 | GC | Microstructure | thickness | syn | precuneus | 0.715016 | 0.249397 | 0.386797 | 543 | (0.7, 1, 0.1) |
| 7 | GC | Cortical expansion | scaling(HCP) | syn | entorhinal | 0.713512 | 0.250491 | 0.318005 | 76 | (0.3, 3, 0.7) |
| 8 | GC | AD-related | *MAPT* | spread | entorhinal | 0.708057 | 0.199068 | 0.486822 | 314 | (0.3, 3, 0.1) |
| 9 | FC | AD-related | *MAPT* | mis | precuneus | 0.707129 | 0.236139 | 0.496083 | 218 | (0.9, 3, 0.3) |
| 10 | FC | Neurotransmission | 5-HT<sub>4</sub> | mis | entorhinal | 0.706228 | 0.218200 | 0.495086 | 252 | (0.9, 3, 0.3) |

Classical models originating from the entorhinal cortex (S1 and S3) or precuneus (S2 and S4) and diffusing through anatomical connections are bolded. The optimal time points (total time = 30,000) are provided, along with the selected hyperparameters: stay probability, transform rate, and velocity.

**Table S6. Performance of models better than null for four tau load subtypes. Related to Fig. 5**.

**(Attached Supplementary_Table6_Subtypes_mdoels_performance.xlsx)**

The optimal time points (total time = 30,000) are provided, along with the selected hyperparameters: stay probability, transform rate, and velocity.

* MSE was calculated using normalized simulated tau values, scaled to match the range of observed tau.

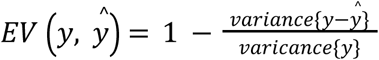

R = Pearson correlation coefficient; EV = explained variance; MSE = mean squared error; syn = synthesis; mis = misfold; clear = clearance; Aβ = β amyloid; SC = structural connectivity; FC = functional connectivity; GC = gene coexpression; NS = neurotransmission markerreceptor similarity; LS = laminar similarity; MC = metabolic covariance; EP = electrophysiological; TS = temporal similarity; CBF: cerebral blood flow; CBV: cerebral blood volume; thickness: cortical thickness; scaling: allometric scalling.

**Table S7.**
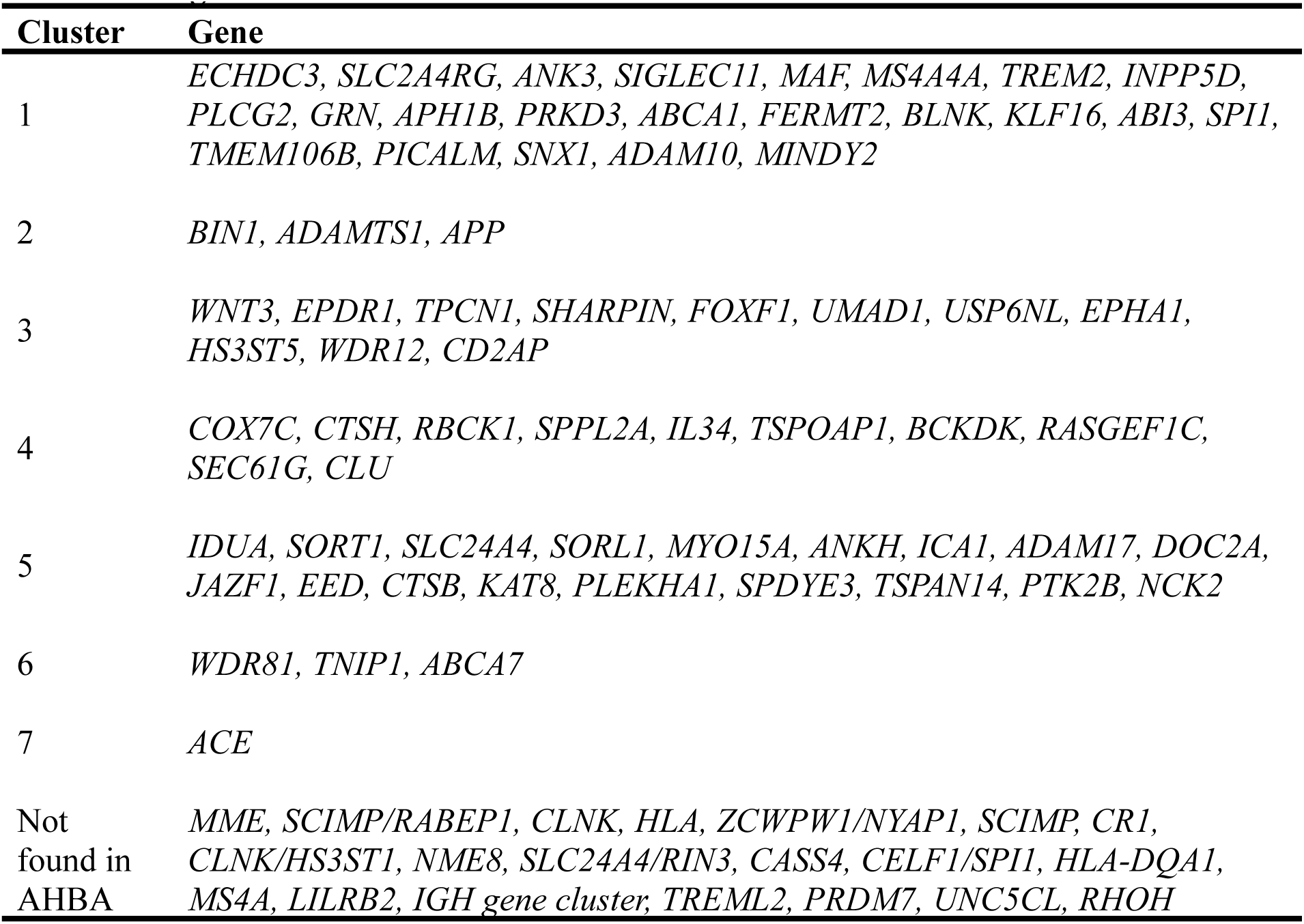
Cluster assginment of AD-related genes identified from three GWAS studies. Related to Fig. S5 and S8.

**Table S8.** Full list of neurotransmission markers and derived ratios. Related to Fig. 4.

| **Abbr.** | **Full name** |
| --- | --- |
| NMDA | **Glutamate** receptor (ionotropic; excitatory) |
| mGluR<sub>5</sub> | Metabotropic **Glutamate** receptor type 5 (metabotropic; excitatory) |
| GABA<sub>A/BZ</sub> | Ionotropic **GABA** receptor type A Benzodiazepine site (ionotropic; inhibitory) |
| M1 | Muscarinic **Acetylcholine** receptor type 1 (metabotropic; excitatory) |
| $\alpha_4\beta_2$ | Alpha-4 beta-2 Nicotinic **Acetylcholine** receptor (ionotropic; excitatory) |
| VAcHT | Vesicular **Acetylcholine** transporter |
| 5-HT<sub>1A</sub> | **Serotonin** receptor type 1A (metabotropic; inhibitory) |
| 5-HT<sub>1B</sub> | **Serotonin** receptor type 1B (metabotropic; inhibitory) |
| 5-HT<sub>2A</sub> | **Serotonin** receptor type 2A (metabotropic; excitatory) |
| 5-HT<sub>4</sub> | **Serotonin** receptor type 4 (metabotropic; excitatory) |
| 5-HT<sub>6</sub> | **Serotonin** receptor type 6 (metabotropic; excitatory) |
| 5-HTT | **Serotonin** transporter |
| NET | **Norepinephrine** transporter |
| D<sub>1</sub> | **Dopamine** receptor type 1 (metabotropic; excitatory) |
| D<sub>2</sub> | **Dopamine** receptor type 2 (metabotropic; inhibitory) |
| DAT | **Dopamine** transporter |
| H<sub>3</sub> | **Histamine** receptor type 3 (metabotropic; inhibitory) |
| MOR | Mu-**Opioid** receptor (metabotropic; inhibitory) |
| CB<sub>1</sub> | **Cannabinoid** receptor type 1 (metabotropic, -) |
| E:I | $(5-HT_{2A}+5-HT_4+5-HT_6+D_1+M_1+\alpha_4\beta_2+NMDA+mGluR_5)/(5-HT_{1A}+5-HT_{1B}+D_2+GABA_{A/BZ}+H_3+MOR)$ |
| E:I<sub>metabotropic</sub> | $(5-HT_{2A}+5-HT_4+5-HT_6+D_1+M_1+mGluR_5)/(5-HT_{1A}+5-HT_{1B}+D_2+H_3+MOR)$ |
| E:I<sub>ionotropic</sub> | $(\alpha_4\beta_2+NMDA)/GABA_{A/BZ}$ |
| E:I<sub>Glu/GABA</sub> | $(NMDA+mGluR_5)/GABA_{A/BZ}$ |
| E:I<sub>Glu/GABA(ion)</sub> | $NMDA/GABA_{A/BZ}$ |
| R/Re<sub>ACh</sub> | $(M_1 + \alpha_4\beta_2)/VAcHT$ |
| NT/NM | $(GABA_{A/BZ}+mGluR_5+NMDA)/(5-HT_{1A}+5-HT_{1B}+5-HT_{2A}+5-HT_4+5-HT_6+\alpha_4\beta_2+D_1+D_2+M_1+H_3+MOR)$ |
| 5-HT<sub>1A</sub>/5-HT<sub>1B</sub> | 5-HT<sub>1A</sub>/5-HT<sub>1B</sub> |
| DAT/NET | DAT/NET |

## Notes

### Summary of Updates

revised version based on reviewers' comments. Changed Fig.1,2 add Fig.6. Changed extended data figures and supplementary

